# Synapse protein signatures in cerebrospinal fluid and plasma predict cognitive maintenance versus decline in Alzheimer’s disease

**DOI:** 10.1101/2024.07.22.604680

**Authors:** Hamilton Se-Hwee Oh, Deniz Yagmur Urey, Linda Karlsson, Zeyu Zhu, Yuanyuan Shen, Amelia Farinas, Jigyasha Timsina, Ian H. Guldner, Nader Morshed, Chengran Yang, Daniel Western, Muhammad Ali, Yann Le Guen, Alexandra Trelle, Sanna-Kaisa Herukka, Tuomas Rauamaa, Mikko Hiltunen, Anssi Lipponen, Antti J. Luikku, Kathleen L. Poston, Elizabeth Mormino, Anthony D. Wagner, Edward N. Wilson, Divya Channappa, Ville Leinonen, Beth Stevens, Alexander J. Ehrenberg, Henrik Zetterberg, David A. Bennett, Nico Franzmeier, Oskar Hansson, Carlos Cruchaga, Tony Wyss-Coray

## Abstract

Rates of cognitive decline in Alzheimer’s disease (AD) are extremely heterogeneous, with ages of symptom onset ranging from age 40-100 years and conversion from mild cognitive impairment to AD dementia taking 2-20 years. Development of biomarkers for amyloid-beta (Aβ) and tau protein aggregates, the hallmark pathologies of AD, have improved patient monitoring/stratification and drug development, but they still only explain 20-40% of the variance in cognitive impairment (CI) in AD. To discover additional molecular drivers and biomarkers of AD dementia, we perform cerebrospinal fluid (CSF) proteomics on 3,416 individuals from six deeply phenotyped prospective AD case-control cohorts. We identify synapse proteins as the strongest correlates of CI, independent of Aβ and tau. Using machine learning we derive the CSF YWHAG:NPTX2 synapse protein ratio, a robust correlate of CI, which explains 27% of the variance in CI beyond CSF PTau181:Aβ42, 10% beyond tau PET, and 50% beyond CSF NfL in Aβ positive individuals. We find YWHAG:NPTX2 also increases with normal aging as early as age 20 and increases at a faster rate in *APOE4* carriers and autosomal dominant-AD mutation carriers. Most notably, YWHAG:NPTX2+ individuals (top 25^th^ percentile) are 15-times (HR=15.4 [10.6-22.2]) more likely to experience cognitive decline over 15 years compared to YWHAG:NPTX2– individuals (bottom 25^th^ percentile), and this rises to 19-times (HR=18.9 [10.83-32.9]) with additional stratification by Aβ and phosphorylated tau status. Lastly, we perform plasma proteomics on 4,245 individuals to develop a plasma-based signature of CI which partly recapitulates CSF YWHAG:NPTX2. Overall, our findings underscore CSF YWHAG:NPTX2 and the corresponding plasma signature as robust prognostic biomarkers for AD onset and progression beyond gold-standard biomarkers of Aβ, tau, and neurodegeneration and implicate synapse dysfunction as a core driver of AD dementia.

## MAIN

Alzheimer’s disease (AD) is the most common age-related neurodegenerative disease characterized by decades long buildup of amyloid-beta (Aβ) plaques and neurofibrillary tau tangles followed by dementia^1^. Rates of cognitive decline in Alzheimer’s disease (AD) are extremely heterogeneous, with ages of AD symptom onset ranging from age 40-100 and conversion from mild cognitive impairment (MCI) to AD dementia taking 2-20 years^2^. While the development of cerebrospinal fluid (CSF) and positron emission tomography (PET) biomarkers of Aβ and tau have begun to untangle this heterogeneity and have thereby improved AD diagnosis, patient stratification, and drug development^3–7^, Aβ and tau still only explain 20-40% of the variance in cognitive impairment (CI) in AD^8–11^ (**Extended Data Fig. 1a)**, suggesting the existence of additional drivers of AD dementia that are not captured by biomarkers of primary AD pathologies Aβ and tau. The prevalence of Aβ-positive (Aβ+) cognitively normal aged individuals further underscores the need for increased understanding of what drives AD dementia versus cognitive resilience^12,13^.

The “A/T/N” (Aβ/tau/neurodegeneration) AD biomarker framework^14^, developed by the National Institute on Aging and the Alzheimer’s Association, has provided a structure to investigate and integrate different AD biomarkers. Among CSF biomarkers, Aβ42 is typically used to define “A” positivity and PTau181 to define “T_1_” (phosphorylated secreted tau) positivity^14^. The CSF PTau181:Aβ42 ratio captures both aspects simultaneously^15,16^. “T_2_” is reserved for emerging biomarkers of fibrillary tau proteinopathy, like CSF pT205, CSF MTBR-243^17^, and tau PET^18^. The “N” category includes Aβ- and tau-independent biomarkers of AD such as neurofilament light (NfL) for axon degeneration, and neurogranin for synapse dysfunction^5^. However, these “N” biomarkers explain only a small additional proportion of variance in CI beyond Aβ and tau^5^.

To discover new robust Aβ- and tau-independent correlates of CI in AD, we perform large-scale proteomics (SomaScan, mass-spectrometry) on the CSF of 3,416 individuals across six deeply phenotyped AD case-control cohorts spanning both sporadic and autosomal dominant AD (ADAD): Stanford (includes Alzheimer’s Disease Research Center (Stanford-ADRC), Stanford Aging and Memory Study (SAMS), and Poston cohort), Knight-ADRC, Alzheimer’s Disease Neuroimaging Initiative (ADNI), Dominantly Inherited Alzheimer’s Network (DIAN), BioFINDER2, and Kuopio University Hospital (**Fig. 1a, Supplementary Table 1**). We integrate these CSF proteomics data with CSF and PET biomarkers of Aβ and tau, clinical diagnosis of cognitive function, age, sex, *APOE4* genotype, and ADAD mutation status, and leverage statistical techniques to derive a robust CSF biomarker of CI that explains CI beyond Aβ and tau. Lastly, we derive a plasma proteomic surrogate of the CSF biomarker of CI based on plasma proteomics (SomaScan) data from 4,525 samples across three independent cohorts: Knight-ADRC, Stanford, Religious Order Study/Memory Aging Project (ROSMAP).

**Figure 1.**
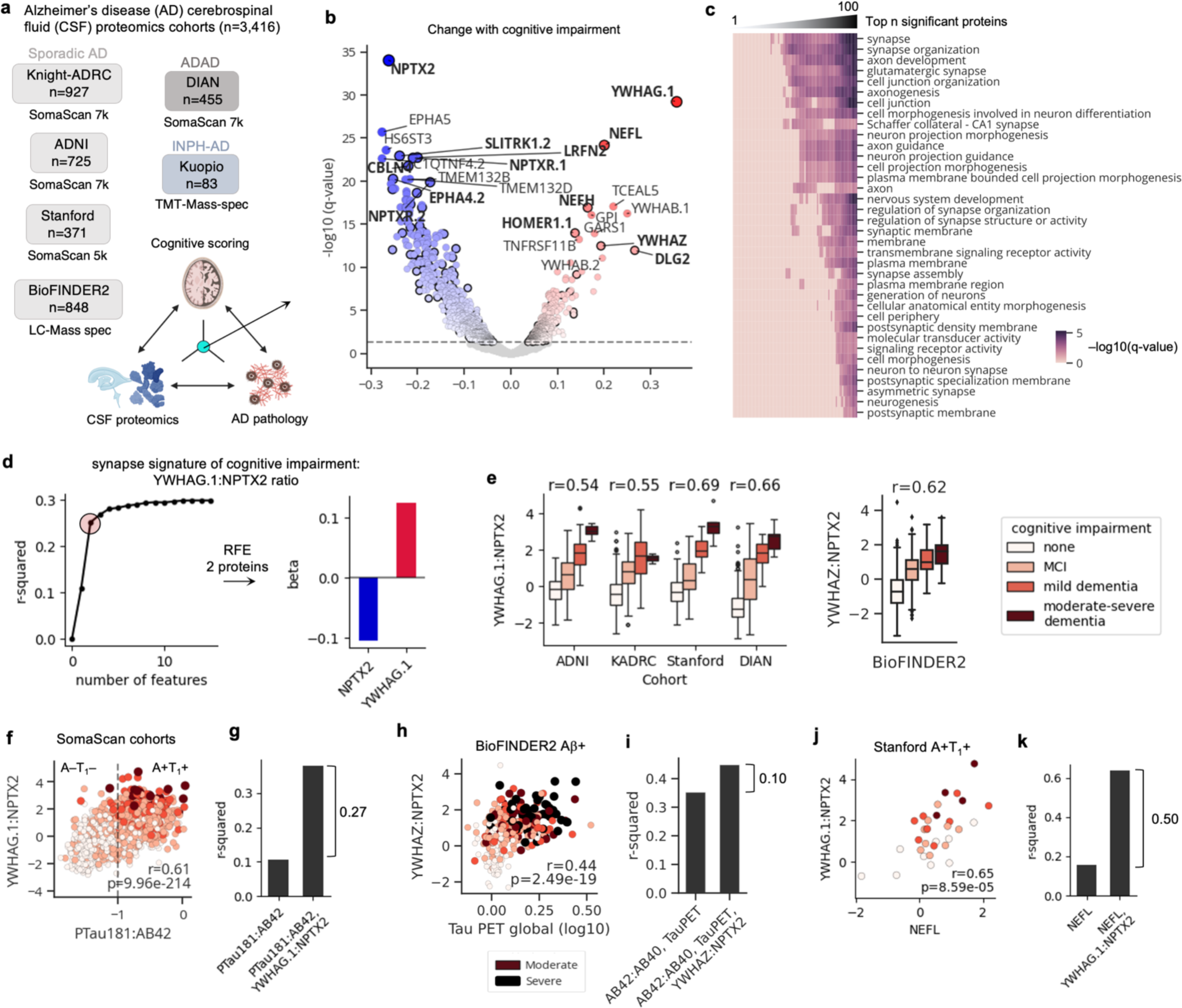
CSF YWHAG:NPTX2 ratio explains a substantial proportion of variance in cognitive impairment beyond amyloid and tau in AD. **a,** Study cohorts. Study design: integration of cerebrospinal fluid proteomics, AD pathology biomarkers, and clinical cognitive scoring to identify molecular correlates of cognitive impairment, independent of AD pathology. **b,** Volcano plot: change with cognitive impairment independent of age, sex, cohort, *APOE4* dose, PTau181:Aβ42, and principal component 1 of the CSF proteome in the Knight-ADRC and ADNI cohorts. Bold indicates synapse proteins based on SynGO database. q-values are Benjamini-Hochberg corrected p-values. **c,** Expanding window pathway enrichment heatmap of all differentially expressed proteins (both up and down). x-axis indicates top-n proteins to include in pathway enrichment test. Cells colored by –log10 (q-value). **d,** A penalized linear model was trained to predict cognitive impairment severity using synapse proteins that significantly change with cognitive impairment. Recursive feature elimination was used to derive a simple model. Scatterplot shows 2 proteins sufficiently capture 83% of the full model performance, determined by cross-validation. Model coefficients show the normalized ratio between YWHAG:NPTX2. **e,** Boxplot showing YWHAG:NPTX2 versus cognitive impairment severity across cohorts with SomaScan data. Boxplot showing YWHAZ:NPTX2 versus cognitive impairment severity based on mass-spectrometry data in BioFINDER2. Standard boxplot metrics used. **f,** Scatterplot showing YWHAG:NPTX2 versus PTau181:Aβ42, colored by cognitive impairment. **g,** R-squared values from multivariate linear models regressing cognitive impairment against covariates displayed on x-axis. The difference between r-squared values between the two models are shown. **h,** Scatterplot showing YWHAZ:NPTX2 versus tau PET in Aβ+ individuals colored by cognitive impairment. **i,** R-squared values from a linear model regression cognitive impairment against covariates displayed on x-axis. The difference between r-squared values between the two models are shown. **h,** Scatterplot showing YWHAG:NPTX2 versus CSF NfL in A+T_1_+ individuals colored by cognitive impairment. **i,** R-squared values from a linear model regression cognitive impairment against covariates displayed on x-axis. The difference between r-squared values between the two models are shown.

### CSF YWHAG:NPTX2 ratio explains a substantial proportion of variance in cognitive impairment beyond amyloid and tau in AD

We performed proteomics on 3,416 CSF samples (3,106 with complete CI diagnosis) from six independent cohorts. To identify CSF proteins that explained additional variance of CI beyond AD pathology, we regressed the global clinical dementia rating (CDR-Global, a clinical cognitive impairment score) against CSF protein levels, while adjusting for CSF PTau181:Aβ42, age, sex, *APOE4*, cohort, and principal component 1 of the proteome (see Methods). We used SomaScan proteomics data with 7,289 protein measurements per sample from the Knight-ADRC and ADNI cohorts for discovery.

We identified 675 significant (Benjamini Hochberg q-value<0.05) upregulated and 721 significant downregulated proteins with CI (**Fig. 1b, Supplementary Table 2-3**). Interestingly, the most significant proteins by q-value were especially enriched at the synapse (based on the SynGO database^19^; **Fig. 1c**). The most upregulated synapse proteins included YWHAG, YWHAZ, YWHAH, NEFL, NEFH, DLG2, HOMER1, MAP1LC3A, PPP3CA, and PPP3R1. The YWHA-family proteins, also referred to as 14-3-3 proteins, are ubiquitously expressed in the body and the CNS and seem to be enriched at neuronal synapses^20,21^. YWHA proteins, DLG2, and calcineurin subunits (PPP3CA and PPP3R1) were especially associated with PTau181:Aβ42^22^ (**Extended Data Fig. 1b-c**), suggesting their changes with CI severity may be co-regulated with Aβ and tau accumulation. In line with these results, previous studies have shown that Aβ42 signaling promotes calcineurin activity^23^ and interestingly, inhibition of calcineurin activity protects mice from Aβ- and tau-induced synapse loss and CI^24,25^. Notably, SMOC1, an extracellular matrix protein previously linked to AD and Aβ plaques^26,27^, was not associated with CI after adjusting for PTau181:Aβ42 (**Extended Data Fig. 1b**), demonstrating the importance of adjusting for PTau181:Aβ42 to identify Aβ- and tau-independent correlates of CI.

The most downregulated proteins with CI included NPTX2, NPTXR, SLITRK1, CBLN4, LRFN2, and EPHA4. These proteins were only weakly negatively associated with PTau181:Aβ42^22^, suggesting their changes with CI are regulated by mechanisms independent of Aβ and tau accumulation (**Extended Data Fig. 1b-c)**. The protein with the strongest decrease was NPTX2, a protein that promotes synaptic plasticity at excitatory synapses^28^ and prevents neuronal network hyperactivity^29^. While it has been studied exclusively in neurons, it is worth noting that the gene is highly expressed in the oligodendrocyte lineage in humans as well^30^. In human brains, NPTX2 mRNA and protein are downregulated in AD neurons based on single-cell RNA-sequencing and immunohistochemistry^31^, suggesting its decrease with CI in CSF may reflect decreased expression in neurons. Interestingly, a recent study showed that overexpression of NPTX2 in the hippocampus of tau-P301S mice protected synapses from complement-mediated glial engulfment^32^.

Given the enrichment of synapse proteins associated with CI, we sought to derive a multi-protein signature that would represent these global changes. Using the ADNI cohort, we trained a penalized linear model to predict CI severity based on levels of 214 synapse proteins that significantly changed with CI in the discovery cohorts. We further used recursive feature elimination (RFE) to simplify the model to facilitate clinical applications (**Fig. 1d**). The model identified the near 1:1 difference between the two most upregulated and downregulated proteins, YWHAG and NPTX2, to be a suitable signature of CI, likely indicative of changes in synapse biology (**Fig. 1d**). Since we log-normalized then z-scored protein levels before analyses, the difference between the normalized protein levels represents a normalized ratio. Notably, ratios between YWHA-family proteins and NPTX2 based on CSF mass spectrometry have previously been shown to be associated with various AD-related phenotypes^33,34^, suggesting reproducibility across cohorts and proteomic platforms. We use the YWHAG:NPTX2 ratio as an indicator of cognitive impairment likely representing changes in synapse biology and, for simplicity, call it a “synapse signature”. Figures refer to YWHAG.1, a specific YWHAG proteoform detected by the Somalogic aptamer (SeqId 4179-57).

We evaluated the associations between YWHAG:NPTX2 and CI across all cohorts with SomaScan data, including the Stanford and DIAN cohorts which were not used for discovery. We found YWHAG:NPTX2 was consistently correlated with CI (ADNI r=0.54, Knight-ADRC r=0.55, Stanford r=0.62, DIAN r=0.66) in all cohorts and in both sporadic AD and ADAD (**Fig. 1e**), confirming YWHAG:NPTX2’s link with the biology of AD dementia. Notably, the correlation between YWHAG:NPTX2 and CI was slightly higher than the correlation between PTau181:Aβ42 and CI across all cohorts (**Extended Data Fig. 1d**).

To determine the robustness of YWHAG:NPTX2 in explaining CI beyond AD pathology, we visualized the relationship between YWHAG:NPTX2 and PTau181:Aβ42, colored by CI severity in a scatterplot (**Fig. 1f**). We observed that while YWHAG:NPTX2 and PTau181:Aβ42 were correlated (r=0.61), low and high levels of YWHAG:NPTX2 further separated A+T_1_+ (log_10_ PTau181:Aβ42 > –1, see methods) individuals into no impairment versus mild-severe dementia, respectively (**Fig. 1f**). Among A+T_1_+ individuals, 62% of individuals with low levels of YWHAG:NPTX2 were cognitively normal and 37% had only MCI, whereas only 4% of individuals with high YWHAG:NPTX2 were cognitively normal and 46% had mild-severe dementia (**Extended Data Fig. 1e**). This pattern was consistent across all cohorts and both sporadic AD and ADAD (**Extended Data Fig. 1e**). Using linear regression, we found that PTau181:Aβ42 explained 10% of the variance in CI in A+T_1_+ individuals, and YWHAG:NPTX2 explained an additional 27%, independent of PTau181:Aβ42 (**Fig. 1h**). YWHAG:NPTX2 was significantly associated with CI independent of PTau181:Aβ42 and age in A+T_1_+ individuals across all cohorts and proteomic platforms (**Extended Data Fig. 1f**). Notably, in the DIAN ADAD-control cohort, we additionally accounted for estimated age of symptom onset, and YWHAG:NPTX2 remained the most significant correlate of CI, demonstrating that it partially explains even the small amounts of heterogeneity in CI in ADAD (**Extended Data Fig. 1f**).

While PTau181:Aβ42 is a robust biomarker of Aβ plaques and phosphorylated secreted tau in the brain, it is not as well correlated with tau tangles, which are known to correlate with CI more strongly^18^. To determine whether YWHAG:NPTX2 explained CI in AD beyond tau tangles (T_2_), we utilized the BioFINDER2 cohort which performed targeted CSF synapse protein mass spectrometry proteomics, tau PET imaging, and measurement of CSF Aβ42:Aβ40. Since YWHAG was not measured, we used YWHAZ, a related protein which was also associated with CI independent of PTau181:Aβ42 albeit not as strongly, based on SomaScan data (**Fig. 1b**). We confirmed YWHAG was highly correlated with YWHAZ (r=0.94), and YWHAG:NPTX2 was highly correlated with YWHAZ:NPTX2 (r=0.94) in the SomaScan cohorts (**Extended Data Fig. 1g**). We visualized the relationship between YWHAZ:NPTX2 and tau PET in Aβ+ individuals, colored by CI severity in a scatterplot (**Fig. 1i**). We observed several interesting patterns. First, we found a moderate correlation between YWHAZ:NPTX2 and tau PET (r=0.44). Second, we observed that all individuals with above moderate levels of tau had above moderate levels of YWHAZ:NPTX2, but not vice versa, suggesting that YWHAZ:NPTX2 may change before tau during AD progression. Third, we observed that YWHAZ:NPTX2 and tau PET independently explained CI severity. Using linear regression, we found that Aβ42:Aβ40 and tau PET together explained 35% of the variance in CI in Aβ+ individuals, and YWHAZ:NPTX2 explained an additional 10%, independent of Aβ42:Aβ40 and tau PET (**Fig. 1j**). YWHAZ:NPTX2’s association with CI was robust to additional adjustment with age, *APOE4* dose, and sex (**Extended Data Fig. 1h**).

We next tested whether YWHAG:NPTX2 explained CI beyond CSF NfL, the current gold-standard neurodegeneration (“N”) biomarker for AD and other neurodegenerative diseases. We measured CSF NfL with Olink proteomics – which is highly concordant with the Simoa assay, r>0.9^35^ – from Stanford participants using the same CSF sample as was analyzed with Somalogic proteomics. Subsetting to the 31 A+T_1_+ individuals with matched proteomic data and CI scoring, we observed that YWHAG:NPTX2 and NfL were correlated (r=0.65), but importantly, YWHAG:NPTX2 explained an additional 50% of the variance in CI beyond NfL (**Fig. 1j-k**).

Together, these results show that CSF synapse proteins, some with established causal roles in synaptic/cognitive resilience to AD pathology in mouse models (i.e. calcineurin, NPTX2), are among the strongest correlates of CI severity independent of Aβ and tau in humans, and that the CSF YWHAG:NPTX2 ratio is a synapse protein signature that explains a major proportion of variance in CI in AD beyond gold-standard biomarkers of Aβ, tau, and neurodegeneration.

### CSF YWHAG:NPTX2 ratio increases with normal aging and pre-symptomatic AD

Since age is the strongest risk factor for AD onset, we wondered whether YWHAG:NPTX2 increases during normal aging before CI. We examined changes in YWHAG:NPTX2 with age in cognitively normal individuals across the human lifespan from all cohorts with SomaScan data. Surprisingly, we found that YWHAG:NPTX2 increased with age not only in later decades, but also in the earliest decades of adulthood, ∼30 years before changes in PTau181:Aβ42 (**Fig. 2a**). This pattern was replicated in the BioFINDER2 study in which proteins were measured with mass spectrometry (**Extended Data Fig. 2a**).

**Figure 2.**
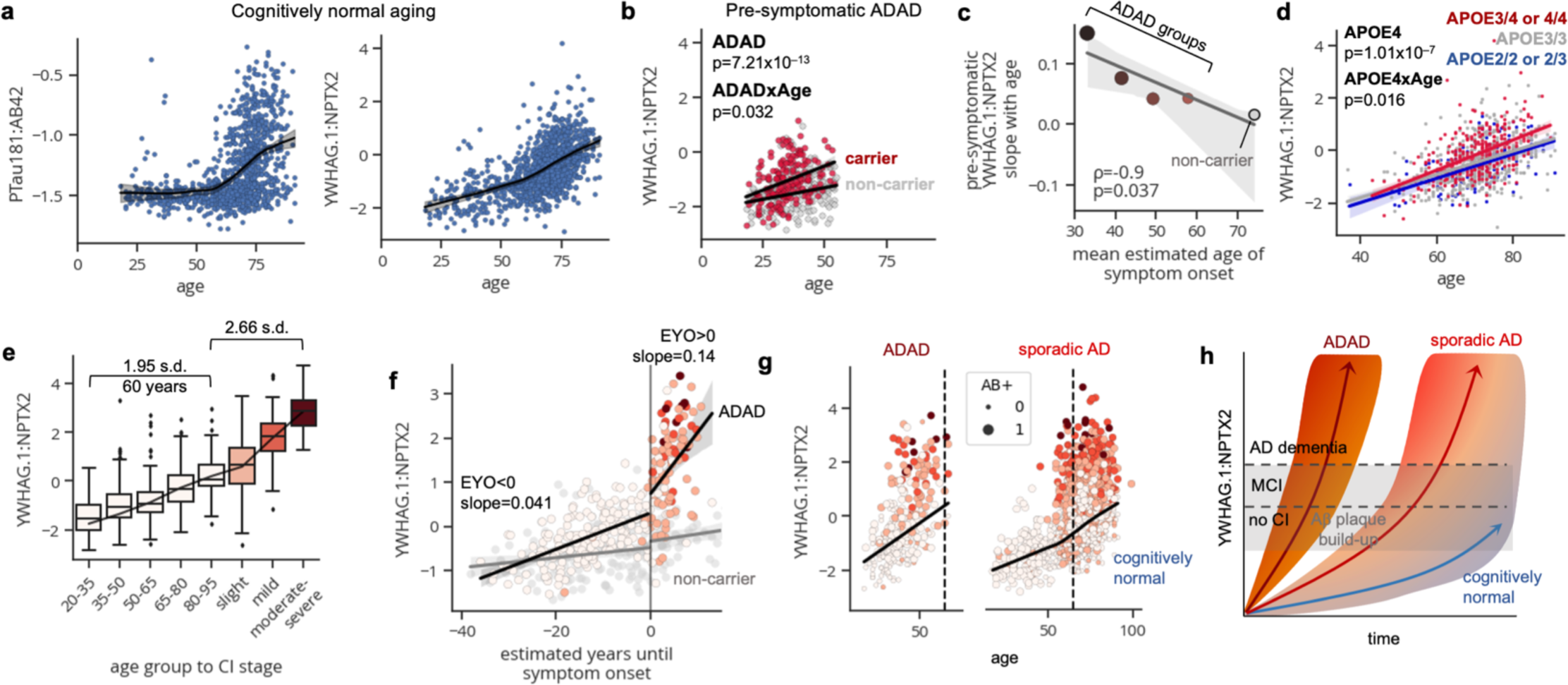
CSF YWHAG:NPTX2 ratio increases with normal aging and pre-symptomatic AD. **a,** Changes with age of YWHAG:NPTX2 and PTau181:Aβ42 in cognitively normal individuals non-ADAD mutation carriers across cohorts with SomaScan data. **b,** Changes with age of YWHAG:NPTX2 in cognitively normal individuals under age 55 stratified by ADAD mutation carrier status. Results from a linear model regressing YWHAG:NPTX2 against ADAD carrier status, age, and their interaction are shown. **c,** Spearman correlation between mean estimated age of onset (EAO) and slope of YWHAG:NPTX2 change with age per EAO-binned ADAD group. Data from non-carriers shown for comparison. **d,** Changes with age of YWHAG:NPTX2 in cognitively normal non-ADAD mutation carriers stratified by *APOE* genotype. Results from a linear model regressing YWHAG:NPTX2 against *APOE4* dose, age, and their interaction are shown. **e,** Changes in YWHAG:NPTX2 across different age groups and CI stages. Standard boxplot metrics used. **f,** Changes with estimated years until symptom onset of YWHAG:NPTX2 stratified by ADAD-carrier status. ADAD-carrier points colored by CI stage. Slopes before and after estimated symptom onset shown. **g,** Changes in YWHAG:NPTX2 with age and CI stages for all individuals shown. Points colored by CI stage and sized by Aβ positivity. **h,** Schematic of proposed model showing that changes in YWHAG:NPTX2 with cognitively normal aging underlie age of AD onset.

To determine whether changes with age in YWHAG:NPTX2 may precede AD symptom onset, we leveraged data from ADAD mutation carriers in the DIAN cohort who have genetically determined earlier onset AD compared to non-carriers. Specifically, we examined whether YWHAG:NPTX2 had a steeper increase with age in pre-symptomatic ADAD carriers compared to non-carriers. We tested a linear model regressing YWHAG:NPTX2 against carrier status, age, and their interaction, among cognitively normal individuals under age 55, the age range where PTau181:Aβ42 levels are normal in non-carrier individuals (**Fig. 2a**). Interestingly, we found that ADAD carriers had significantly higher YWHAG:NPTX2 (p= 7.21×10^−13^) and a steeper increase in YWHAG:NPTX2 with age (interaction p=0.032) during this cognitively normal phase (**Fig. 2b**). Visualizing the differences in slopes with age, we observed that ADAD carriers had double the slope of YWHAG:NPTX2 compared to non-carriers (**Fig. 2b**).

To explore this phenomenon further, we leveraged the fact that ADAD mutations have varying degrees of severity, with estimated ages of symptom onset ranging from age 25-65 depending on the mutation^27,36^. To determine whether age-related slopes of YWHAG:NPTX2 scaled with ADAD mutation severity, we grouped pre-symptomatic carriers into different bins based on estimated age of onset (<35, 35-45, 45-55, 55-65) and calculated the age-related YWHAG:NPTX2 slopes per bin. We tested the correlation between the mean estimated age of symptom onset per bin with their respective YWHAG:NPTX2 slopes and observed a strong negative correlation (Spearman r=–0.9, p=0.037), whereby those with earlier ages of symptom onset had steeper increases in YWHAG:NPTX2 during the pre-symptomatic phase (**Fig. 2c**, **Extended Data Fig. 2b**).

We then examined the effects of *APOE* genotype, the leading genetic risk factor for sporadic AD, on YWHAG:NPTX2 aging slopes. We tested a linear model regressing YWHAG:NPTX2 against *APOE4* dose, age, and their interaction in cognitively normal individuals across the lifespan from the Knight-ADRC, ADNI, and Stanford SomaScan cohorts. Like ADAD mutation carrier status, *APOE4* was significantly associated with higher YWHAG:NPTX2 (p= 1.01×10^−7^) and a steeper increase in YWHAG:NPTX2 with age (p=0.016). Visualizing the differences in slopes between *APOE4* carriers, *APOE*3/3 homozygotes, and *APOE*2 carriers, we observed that *APOE4* carriers had a 33% steeper increase in YWHAG:NPTX2 compared to *APOE*3/3 homozygotes (**Fig. 2d**), in line with the earlier age of onset in *APOE4* carriers. *APOE*2 carriers did not have a significantly different increase in YWHAG:NPTX2 compared to *APOE*3/3 homozygotes, though we suspect this may be due to limited sample size.

Our analyses thus far revealed that YWHAG:NPTX2 increases with both normal aging (**Fig. 2a-d**), pre-symptomatic AD, as well as CI severity during AD progression (**Fig. 1d-k**). We next sought to compare the degrees to which YWHAG:NPTX2 increases during these two stages.

We compared YWHAG:NPTX2 changes with normal aging – 20-year binned age groups from young (age 20-35) to old (age 80-95) – and with AD progression – cognitively normal old (age 80-95) to moderate-severe CI (age 65-95). We found that YWHAG:NPTX2 increased by 1.9 standard deviations over 60 years of normal aging and then 2.7 standard deviations from aged to dementia (**Fig. 2e**). Though there is no age difference between cognitively normal versus dementia in our cohorts since they are age-matched case-control studies, assuming a typical ∼20 years of time for progression from Aβ+ cognitively normal to AD dementia based on population-based studies^37,38^, our data suggest that AD progression recapitulates ∼84 years of “normal” age-related increases in YWHAG:NPTX2, representing a stark ∼4.3x increase in its slope during AD progression compared to normal aging.

We examined this phenomenon in ADAD by plotting the change in YWHAG:NPTX2 across estimated time until symptom onset. We compared YWHAG:NPTX2 slopes before and after estimated symptom onset and, similar to our estimates in sporadic AD, we observed a 3.4x increase in the YWHAG:NPTX2 slope during ADAD symptom progression compared to the pre-symptomatic phase (**Fig. 2f**). Notably, YWHAG:NPTX2 increased in ADAD ∼20 years before estimated symptom onset compared to non-carriers.

To obtain a birds-eye view of the relationship between age- and dementia-related changes in YWHAG:NPTX2 across sporadic AD and ADAD, we plotted all individuals on a scatterplot, showing YWHAG:NPTX2 versus age, colored by CI stage and sized by amyloid positivity (**Fig. 2g**). We confirmed the extremely accelerated increase in YWHAG:NPTX2 among ADAD mutation carriers leading to early onset AD, as well as the widespread heterogeneity in non-carriers leading to sporadic AD in some and cognitive maintenance in others, despite amyloid positivity and old age (**Fig. 2g-h**).

Collectively, these results demonstrate that YWHAG:NPTX2, a robust correlate of CI severity in AD, increases substantially with cognitively normal aging and pre-symptomatic AD.

### CSF YWHAG:NPTX2 ratio predicts future tau accumulation and cognitive decline beyond Aβ and tau

We next sought to determine the potential clinical utility of YWHAG:NPTX2 in predicting future AD onset and progression. First, we leveraged Aβ and tau PET imaging data that were collected 4-15 years after CSF draw in the ADNI cohort to determine whether YWHAG:NPTX2 could predict future Aβ-driven tau accumulation (**Fig. 3a**). Using linear regression, we found that YWHAG:NPTX2 modified the future association between Aβ and tau PET (YWHAG:NPTX2 x AβPET interaction p=6.84×10^−4^), while adjusting for baseline CI, PTau181:Aβ42, age, sex, and *APOE4*. Among individuals with high future Aβ load, high baseline YWHAG:NPTX2 was associated with higher future tau-PET, while low YWHAG:NPTX2 was associated with limited Aβ-related tau-PET increase (**Fig. 3b**). These results align with previous studies which show that Aβ combined with synapse dysfunction and neuronal hyperactivity drives tau accumulation and propagation^39,40^ and additional studies which show that CSF levels of synaptic protein GAP43 modifies the rate of Aβ-driven tau accumulation^41^.

**Figure 3.**
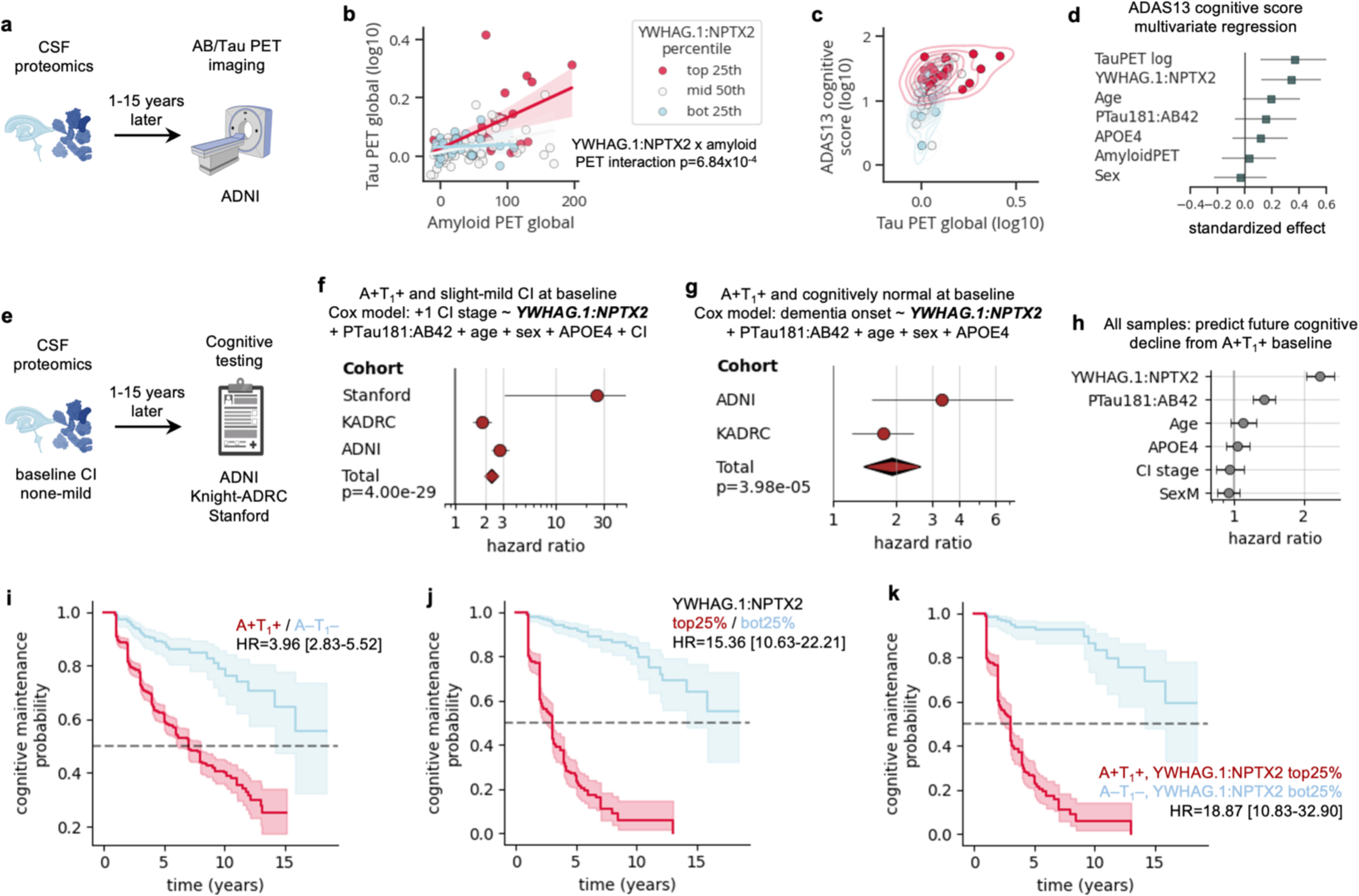
CSF YWHAG:NPTX2 ratio predicts future tau accumulation and cognitive decline beyond Aβ and tau. **a,** ADNI cohort analyses for **b-d**, correlating baseline CSF YWHAG:NPTX2 and PTau181:Aβ42 with future amyloid and tau PET imaging data and cognitive scoring data. **b,** Scatterplot showing future tau PET (global SUVR) versus amyloid PET (global centiloid), colored by percentiles of YWHAG:NPTX2. **c,** Scatterplot showing future ADAS13 cognitive score versus tau PET (global SUVR) colored by percentiles of YWHAG:NPTX2 in Aβ+ individuals. **d,** Results from a multivariate linear mode regressing ADAS13 cognitive score against YWHAG:NPTX2, tau PET, age, *APOE4*, PTau181:Aβ42, sex, and Aβ PET, in Aβ+ individuals. Standardized effects and 95% confidence intervals shown. **e,** ADNI, Knight-ADRC, and Stanford analyses for **f-k**, associating baseline CSF YWHAG:NPTX2 with future cognitive decline. **f,** Cox proportional hazard regression was used to associate YWHAG:NPTX2 with future cognitive decline (defined by a stepwise increase in cognitive impairment stage) in A+T_1_+ individuals with MCI-mild dementia, while adjusting for PTau181:Aβ42, *APOE4*, age, sex, and cognitive impairment stage. Results from a cross-cohort meta-analysis shown. Hazard ratios and 95% confidence intervals shown. **g,** As in **f**, but for predicting dementia onset in A+T_1_+ cognitively normal individuals. **h,** Cox proportional hazard regression was used to associate YWHAG:NPTX2 with future cognitive decline (defined by a stepwise increase in cognitive impairment stage) in all A+T_1_+ individuals across all cohorts, while adjusting for PTau181:Aβ42, *APOE4*, age, sex, and cognitive impairment stage. Hazard ratios and 95% confidence intervals for each covariate shown. **i,** Kaplan Meier curve showing rates of future cognitive decline (defined by a stepwise increase in cognitive impairment stage), in A+T_1_+ versus A–T_1_– individuals. Hazard ratio and 95% confidence interval shown. **j,** As in **i**, but for YWHAG:NPTX2+ (top 25^th^ percentile) versus YWHAG:NPTX2– (bottom 25^th^ percentile) individuals. **k,** As in **i**, but for A+T_1_+YWHAG:NPTX2+ versus A–T_1_–YWHAG:NPTX2– individuals.

More important than predicting future tau tangle build-up is predicting future cognitive decline. Thus, we visualized the relationship between future ADAS13 cognitive score, future tau load, and baseline YWHAG:NPTX2 in future Aβ PET positive individuals on a scatterplot (**Fig. 3c**). We used the ADAS13 score as it is more sensitive and has a higher dynamic range than CDR-Global. Interestingly, we observed that among individuals with low to mild tau build-up, YWHAG:NPTX2 distinguished cognitively normal versus impaired individuals (**Fig. 3c**). All individuals with high tau PET had high YWHAG:NPTX2. We confirmed YWHAG:NPTX2 was significantly associated with future cognitive decline, while adjusting for tau tangle load and several additional covariates (**Fig. 3d**).

To more broadly assess whether YWHAG:NPTX2 could predict future cognitive decline independent of Aβ and tau, we used data from all cohorts with longitudinal cognitive follow-up (ADNI, Knight-ADRC, Stanford; n=1,365 subjects; **Fig. 3e**). We used the CDR-Global CI staging as this was commonly measured across all cohorts. We analyzed both dementia progression from a MCI-mild dementia baseline, as well as dementia onset from a cognitively normal baseline.

First, we employed Cox proportional hazard regression to test the association between YWHAG:NPTX2 and a future increase in CI stage among A+T_1_+ individuals with MCI-mild dementia over 1-15 years, while adjusting for baseline CI, PTau181:Aβ42, age, sex, and *APOE4* dose (**Fig. 3f, Supplementary Table 4**) in each cohort. YWHAG:NPTX2 significantly predicted future cognitive decline across all cohorts, and in a cross-cohort meta-analysis, a standard deviation increase in YWHAG:NPTX2 conferred a 134% increase in risk of cognitive decline (meta hazard ratio=2.34, meta p=3.99×10^−29^; **Fig. 3f**).

We then investigated whether YWHAG:NPTX2 could predict dementia onset from A+T_1_+ cognitively normal individuals (Stanford cohort was not included due to low event sample size). We found that YWHAG:NPTX2 significantly predicted dementia onset across all cohorts, and in a cross-cohort meta-analysis, a standard deviation increase in YWHAG:NPTX2 conferred a 92% increase in risk of conversion from cognitively normal to dementia, while adjusting for PTau181:Aβ42, age, sex, and *APOE4* dose (meta hazard ratio=1.92, meta p=4.00×10^−5^; **Fig. 3g, Supplementary Table 5**).

Given the similar hazard ratios across CI stages, we aggregated data from all A+T_1_+ individuals who had either no CI, MCI, or mild dementia across cohorts and found that YWHAG:NPTX2 was by far the strongest predictor of future cognitive decline among covariates (hazard ratio=2.35, p=2.28×10^−35^; **Fig. 3h**).

Like with AT_1_ status, binning individuals into binary +/– groups provides a simple framework that can aid in patient stratification. Thus, we stratified patients into YWHAG:NPTX2+/– groups based on the upper and lower 25^th^ percentiles, and tested the ability of AT_1_ status and YWHAG:NPTX2 status in predicting future cognitive decline, individually and together. As done previously, we aggregated data from all individuals who had either no CI, MCI, or mild dementia across cohorts. Based on AT_1_ status alone, we found that A+T_1_+ individuals had a roughly 4-times increased risk of future cognitive decline compared to A–T_1_– individuals (hazard ratio=3.96, p=5.94×10^−16^; **Fig. 3i**). Surprisingly, based on YWHAG:NPTX2 status alone, YWHAG:NPTX2+ individuals had a striking ∼15-times increased risk of future cognitive decline compared YWHAG:NPTX2– individuals (hazard ratio=15.36, p=8.04×10^−48^; **Fig. 3j**). Combining the two biomarkers led to even stronger predictions, as A+T_1_+YWHAG:NPTX2+ individuals had a nearly 19-times increased risk of future cognitive decline compared to A–T_1_–YWHAG:NPTX2– individuals (hazard ratio=18.87, p=3.74×10^−25^; **Fig. 3k**). No additional covariates were included in these Cox models, demonstrating the power of these biomarkers alone in predicting future cognitive decline versus maintenance.

Together, these results demonstrate that YWHAG:NPTX2 provides additional prognostic clinical utility beyond gold standard AD biomarkers.

### Plasma proteomic signature of cognitive impairment partly recapitulates CSF YWHAG:NPTX2 ratio, predicting AD onset and progression

While CSF biomarkers provide important insights for AD research and clinical trials, the invasiveness of CSF extraction limits their prognostic utility and widespread clinical use. Thus, we sought to derive a plasma proteomics-based biomarker of CI that could recapitulate CSF YWHAG:NPTX2. We performed SomaScan plasma proteomics on 4,245 samples from the Knight-ADRC, Stanford, and Religious Order Study/Memory Aging Project (ROSMAP) cohorts (**Fig. 4a, Supplementary Table 6**). 3,899 samples had complete CI diagnosis, and 503 samples from the Knight-ADRC and Stanford cohorts were collected within 6 months of CSF samples from the same individuals, which enabled us to directly correlate plasma and CSF protein signatures.

**Figure 4.**
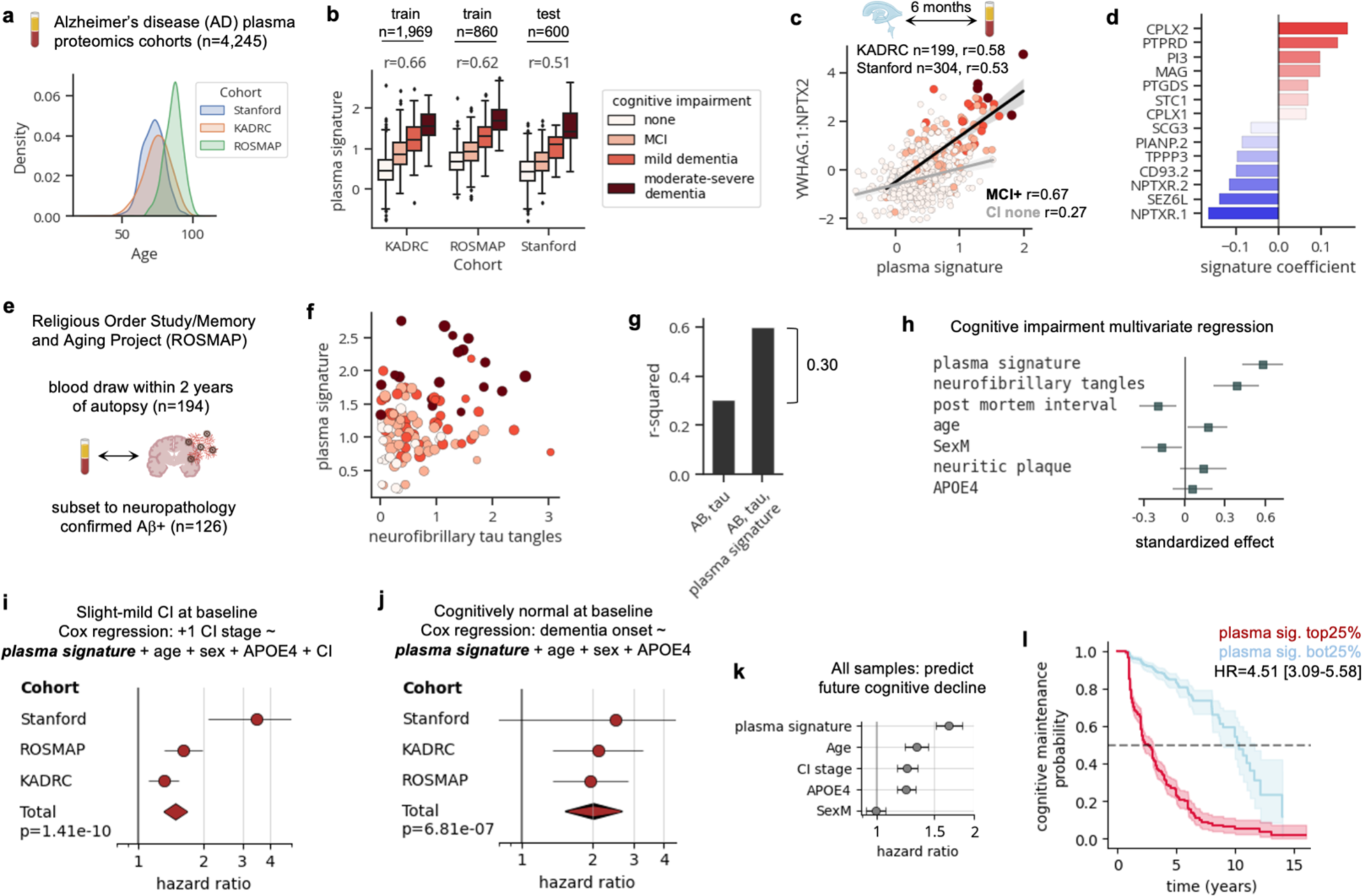
Plasma proteomic signature of cognitive impairment partly recapitulates CSF YWHAG:NPTX2 ratio, predicting AD onset and progression. **a**, SomaScan plasma proteomics data were collected from Stanford, Knight-ADRC, and ROSMAP cohorts. A plasma protein-based signature of cognitive impairment was trained using data from Knight-ADRC and ROSMAP and tested on Stanford. **b,** Boxplot showing plasma signature of cognitive impairment versus actual cognitive impairment severity across cohorts. Pearson correlations shown. **c,** Scatterplot showing correlation between plasma signature and CSF YWHAG:NPTX2 in Knight-ADRC and Stanford cohorts. Only patients (n=503) for which CSF and plasma were collected within 6 months were assessed. Pearson correlations shown. **d,** Bar plot showing protein coefficients for plasma signature. **e,** ROSMAP cohort was used to associate the plasma signature with cognitive impairment, independent of AD neuropathology. A subset of 126 patient plasma samples within 2 years of autopsy and neuropathology confirmed Aβ positivity was assessed. **f,** Scatterplot showing plasma signature versus neurofibrillary tau tangle load, colored by cognitive impairment. **g,** R-squared values from multivariate linear models regressing cognitive impairment against covariates displayed on x-axis. **h,** Results from a multivariate linear model regressing cognitive impairment against the displayed covariates. Standardized effects and 95% confidence intervals shown. **i,** Cox proportional hazard regression was used to associate the plasma signature with future cognitive decline (defined by a stepwise increase in cognitive impairment stage) in individuals with MCI-mild dementia, while adjusting for *APOE4*, age, sex, and cognitive impairment stage. Results from a cross-cohort meta-analysis shown. Hazard ratios and 95% confidence intervals shown. **j,** As in **i**, but for predicting dementia onset in cognitively normal individuals. **k,** Cox proportional hazard regression was used to associate the plasma signature with future cognitive decline (defined by a stepwise increase in cognitive impairment stage) in all individuals across all cohorts, while adjusting for PTau181:Aβ42, *APOE4*, age, sex, and cognitive impairment stage. Hazard ratios and 95% confidence intervals for each covariate shown. **i,** Kaplan Meier curve showing rates of future cognitive decline (defined by a stepwise increase in cognitive impairment stage), in plasma signature+ (top 25^th^ percentile) versus plasma signature– (bottom 25^th^ percentile) individuals. Hazard ratio and 95% confidence interval shown.

We first tested the association between plasma YWHAG:NPTX2 with CI and CSF YWHAG:NPTX2 and found no significant correlations. We then systematically tested several frameworks to optimize correlations between the plasma signature with CI and CSF YWHAG:NPTX2 (see Methods). Briefly, the optimal framework worked as follows: we trained a penalized linear model to predict CI based on a subset of plasma proteins that were 1) enriched for synapse proteins that changed with CI in CSF, 2) not subject to cohort effects and 3) not subject to *APOE* genotype-based proteoform-aptamer binding alterations. We trained the plasma signature of CI in the Knight-ADRC and ROSMAP cohorts and tested in the Stanford cohort.

The plasma signature was correlated with CI across train and test cohorts (**Fig. 4b**; Knight-ADRC r=0.66, ROSMAP r=0.62, Stanford r=0.51). The plasma signature was correlated with CSF YWHAG:NPTX2 (**Fig. 4c**; Knight-ADRC r=0.58, Stanford r=0.53), with stronger correlations observed in individuals with some degree of CI (**Fig. 4c**; CI>=MCI r=0.66, CI=none r=0.28). The proteins with the strongest weights in the plasma signature included CPLX2, PTPRD, PI3, MAG, and PTGDS which increased with CI and NPTXR, SEZ6L, CD93, TPPP3, and PIANP which decreased with CI (**Supplementary Table 7)**. Notably, CPLX2, PTPRD, NPTXR (the receptor for NPTX2), and SEZ6L are synaptic proteins, confirming synapse protein associations with CI across both CSF and plasma (**Fig. 4d**).

To determine whether the plasma signature, like CSF YWHAG:NPTX2, explained CI beyond Aβ and tau in AD we utilized data from the ROSMAP cohort, where comprehensive neuropathological and cognitive phenotyping were performed across most samples. We analyzed a subset of 126 individuals whose blood draws were within 2 years of death and autopsies confirmed neuropathologic diagnosis of AD (neuritic plaques CERAD score=probable or definite; **Fig. 4e**). We visualized the relationship between the plasma signature and the abundance of neurofibrillary tau tangles, colored by CI severity in a scatterplot (**Fig. 4f**). Strikingly, we observed that high plasma signature levels were correlated with CI beyond tau levels (**Fig. 4f**). Using linear regression, we determined that the plasma signature explained an equal and independent proportion of variance in CI (30%) compared to neuritic Aβ plaque and tau tangle load (30%), which together captured 60% of the variance in CI (**Fig. 4g**). The association between the plasma signature and CI remained robust to additional adjustment with age, sex, *APOE4* dose, and post-mortem interval (**Fig. 4h**).

We then investigated whether the plasma signature could be used to predict future cognitive decline, similar to CSF YWHAG:NPTX2. For each cohort, we employed a Cox proportional hazard regression model to test the association between the plasma signature and a future increase in CI stage over 1-15 years among individuals with MCI-mild dementia, while adjusting for baseline CI, age, sex, and *APOE4* dose (**Fig. 4i, Supplementary Table 8**). The plasma signature significantly predicted future cognitive decline across all cohorts, and in a cross-cohort meta-analysis, a standard deviation increase in the plasma signature conferred a 49% increase in risk of cognitive decline (meta hazard ratio=1.49, meta p=1.41×10^−10^; **Fig. 4i**). We also tested associations with future conversion from cognitively normal to dementia and found a robust association such that in a cross-cohort meta-analysis, a standard deviation increase in the plasma signature conferred a 103% increase in risk of conversion from cognitively normal to dementia, while adjusting for age, sex, and *APOE4* dose (meta hazard ratio=2.03, meta p=6.81×10^−7^; **Fig. 4j, Supplementary Table 9**). After aggregating data from all individuals across cohorts we found that the plasma signature was among the strongest predictors of future cognitive decline among covariates (hazard ratio=1.67, p=3.77×10^−27^; **Fig. 4k**), with *APOE4* dose and baseline CI also having similar effect sizes. We then defined binary +/– groups based on the upper and lower 25^th^ percentiles based on all individuals across cohorts, as done with YWHAG:NPTX2. We found that plasma signature+ individuals had a 4.5 times increase risk of future cognitive decline compared to plasma signature– individuals (hazard ratio=4.51, p=3.01×10^−21^; **Fig. 4l**), with no additional covariate adjustment.

Together, these data show plasma proteomics combined with machine learning can be used to derive a plasma-based protein signature which predicts AD dementia independent of Aβ and tau and partly recapitulates CSF YWHAG:NPTX2.

## DISCUSSION

Overall, our findings reveal that synapse proteins in the CSF and plasma are among the strongest Aβ- and tau-independent correlates of CI in AD, and that from these synapse proteins emerges the CSF YWHAG:NPTX2 ratio, a sparse and robust correlate of CI. We find that YWHAG:NPTX2 increases with cognitively normal aging and predicts AD onset and progression in both sporadic AD and ADAD across 6 independent deeply phenotyped AD cohorts, indicating that YWHAG:NPTX2 represents a biological process that is central to AD dementia. Most notably, YWHAG:NPTX2+ individuals are 15-times more likely to experience cognitive decline over 15 years compared to YWHAG:NPTX2– individuals, and this rises to 19-times with additional stratification by AT_1_ status.

What the levels of YWHAG:NPTX2 in CSF precisely represent is unclear. Based on the literature, we speculate that it reflects aspects of synapse dysfunction and neuronal hyperactivity-induced synapse loss. Neuronal pentraxins (i.e. NPTX2, NPTXR, NPTX1) have been previously proposed as biomarkers of synaptic activity^42,43^ as NPTX2^-/-^/NPTXR^-/-^ mice have major GluA4 loss and network hyperactivity^29^. Furthermore, NPTX2^-/-^ mice display increased complement mediated microglial engulfment of synapses, and overexpression of NPTX2 in tau-P301S mice protects synapses from complement mediated microglial engulfment^32^. Though the roles of YWHAG in the brain are less understood, it along with other YWHA-family proteins have been shown to be localized at synapses^20^, and mutations in YWHAG cause childhood epilepsy^44^. YWHAG also binds to phosphorylated tau^45^ and phosphatidyl-serine^20^ which is involved in synaptic pruning^46^.

In addition to reported roles of YWHAG and NPTX2 in synapse biology, our study shows CSF YWHAG:NPTX2 is associated with various aspects of AD including CI, normal aging, genetically driven Aβ overproduction (ADAD), and tau accumulation, which together strongly implicate its relevance to synapse dysfunction. To elaborate, similar to YWHAG:NPTX2 (**Fig. 1f-k**), synapse loss is the most robust histological correlate of CI, beyond Aβ and tau^47^. Second, synapse dysfunction and loss, rather than overt neuron loss, is a major hallmark of mammalian brain aging that is closely linked with cognitive decline in non-human primates^48^ (**Fig. 2a**). Third, Aβ oligomers cause synapse loss and neuronal hyperactivity^47^, akin to how ADAD mutations – which presumably lead to Aβ overproduction – are associated with a faster increase in YWHAG:NPTX2 with age (**Fig. 2b-d**). Lastly, neuronal hyperactivity enhances tau propagation^40^, which aligns with the positive association between YWHAG:NPTX2 and future tau PET (**Fig. 1i**, **Fig. 3b**).

Together, these data suggest CSF YWHAG:NPTX2 is likely a measure of synapse dysfunction and point to synapse dysfunction as a promising therapeutic target to promote cognitive resilience in the presence of Aβ and tau. Perhaps therapies that restore age- and AD-related loss of NPTX2 expression in excitatory neurons, such as NPTX2 gene therapy or delivery of NPTX2 activators (i.e. brain derived neural growth factor, BDNF), could prevent synapse loss and cognitive decline, with CSF YWHAG:NPTX2 as a primary endpoint in such clinical trials. Future studies should determine whether CSF YWHAG:NPTX2 is correlated with CI and synapse dysfunction in non-AD dementias including frontotemporal dementia (FTD) and amyotrophic lateral sclerosis (ALS).

Beyond biological and therapeutic implications, we show comprehensive evidence that CSF YWHAG:NPTX2 would provide major additional diagnostic and prognostic utility as an AD biomarker under the “N” category of the A/T/N framework. We further show the development of a plasma proteomic signature of CI that partly recapitulates the characteristics of CSF YWHAG:NPTX2. Notably, the highest weighted proteins in the plasma signature are synapse proteins that have been previously identified as brain-specific proteins linked to brain aging^49^. Future advances in proteomics and machine learning frameworks will lead to sparse, scalable CSF/plasma biomarkers of synapse dysfunction to be used broadly for patient monitoring, clinical trials, and research.

## METHODS

### PARTICIPANTS

#### Stanford (ADRC, SAMS, BPD, SCMD)

Plasma and CSF collection, processing, and storage for all Stanford cohorts were performed using a single standard operating procedure. All studies were approved by the Institutional Review Board of Stanford University and written informed consent or assent was obtained from all participants or their legally authorized representative.

Blood collection and processing were done according to a rigorous standardized protocol to minimize variation associated with blood draw and blood processing. Briefly, about 10 cc of whole blood was collected in 4 vacutainer ethylenediaminetetraacetic acid (EDTA) tubes (Becton Dickinson vacutainer EDTA tube) and spun at 1800 x g for 10 mins to separate out plasma, leaving 1 cm of plasma above the buffy coat and taking care not to disturb the buffy coat to circumvent cell contamination. Plasma was aliquoted into polypropylene tubes and stored at - 80°C. Plasma processing times averaged approximately one hour from the time of the blood draw to the time of freezing and storage. All blood draws were done in the morning to minimize the impact of circadian rhythm on protein concentrations.

CSF was collected via lumbar puncture using a 20-22 G spinal needle that was inserted in the L4-L5 or L5-S1 interspace. CSF samples were immediately centrifuged at 500 x g for 10 mins, aliquoted in polypropylene tubes and stored at -80°C.

Plasma samples from all Stanford cohorts were sent for proteomics using the SOMAscan platform (SOMAscan7k) in the same batch. CSF samples from all Stanford cohorts were sent for proteomics using the SOMAscan platform (SOMAscan5k) in the same batch. Core CSF AD biomarkers Aβ_42_, Aβ_40_, and pTau181 were measured using the fully automated Lumipulse *G* 1200 instrument (Fujirebio US, Malvern, PA) as previously described^50,51^ for all Stanford cohorts. Descriptions for each cohort are provided below.

A total of 1160 plasma samples (738 participants, longitudinal sampling) and 371 CSF samples (371 participants, 1 sample from each) from Stanford were included in this study. Per-cohort sample sizes are as follows: ADRC plasma n=827 (423 participants), CSF n=113; SAMS plasma n=222 (215 participants), CSF n=169. BPD plasma n=55 (55 participants), CSF n=68. SCMD plasma n=45 (45 participants), CSF n=21.

#### Stanford-Alzheimer’s Disease Research Center (ADRC)

Samples were acquired through the National Institute on Aging (NIA)-funded Stanford Alzheimer’s Disease Research Center (Stanford-ADRC). The Stanford-ADRC cohort is a longitudinal observational study of clinical dementia subjects and age-sex-matched nondemented subjects. All healthy control participants were deemed cognitively unimpaired during a clinical consensus conference that included board-certified neurologists and neuropsychologists. Cognitively impaired subjects underwent Clinical Dementia Rating and standardized neurological and neuropsychological assessments to determine cognitive and diagnostic status, including procedures of the National Alzheimer’s Coordinating Center (https://naccdata.org/). Cognitive status was determined in a clinical consensus conference that included neurologists and neuropsychologists. All participants were free from acute infectious diseases and in good physical condition.

#### Stanford Aging and Memory Study (SAMS)

SAMS is an ongoing longitudinal study of healthy aging. Blood and CSF collection and processing were done by the same team and using the same protocol as in Stanford-ADRC. Neurological and neuropsychological assessments were performed by the same team and using the same protocol as in Stanford-ADRC. All SAMS participants had CDR = 0 and a neuropsychological test score within the normal range; all SAMS participants were deemed cognitively unimpaired during a clinical consensus conference that included neurologists and neuropsychologists.

#### Stanford Biomarkers in PD Study (BPD)

The BPD cohort^52^ was a Michael J Fox Foundation for Parkinson’s Research (MJFF) funded longitudinal study of biological markers associated with cognitive decline in people with a diagnosis of Parkinson’s disease (PD). Research participants were recruited from the Stanford Movement Disorders Center between 2011-2015 with a diagnosis of PD according to UK Brain Bank criteria and required bradykinesia with muscle rigidity and/or rest tremor. All participants completed baseline cognitive, motor, neuropsychologic, imaging, and biomarkers assessments (plasma and optional CSF) including Movement Disorders Society-revised Unified Parkinson’s Disease Rating Scale (MDS-UPDRS). Age-match Healthy Controls (HC) were also recruited to control for age-associated biomarker changes. After comprehensive neuropsychological battery all participants were given a cognitive diagnosis of no cognitive impairment, mild cognitive impairment, or dementia, according to published criteria.

#### Stanford Center for Memory Disorders Cohort Study (SCMD)

The SCMD was an NIA-funded cross-sectional study of people across the cognitive continuum. Participants with mild dementia due to Alzheimer’s (AD) and amnestic mild cognitive impairment (aMCI) were recruited from the Stanford Center for Memory Disorders between 2011-2015. Participants were included if they had a diagnosis of probable AD dementia (amnestic presentation) according to the National Institute on Aging–Alzheimer’s Associationg^53^ (NIA-AA) criteria and a Clinical Dementia Rating (CDR) score of 0.5 or 1, or a diagnosis of MCI according to the NIA-AA criteria^53^, a score of 1.5 SDs below age-adjusted normative means on at least one test of episodic memory, and a CDR score of less than 1. Healthy older controls (HC) were recruited from the community, were selected to have a similar average age as enrolled patients, and were required to have normal neuropsychological performance and CDR of 0. Participants completed cognitive, neuropsychologic, imaging, and biomarker assessments with plasma.

#### Knight-Alzheimer’s Disease Research Center (ADRC)

The Knight-ADRC cohort is an NIA-funded longitudinal observational study of clinical dementia subjects and age-matched controls. Research participants at the Knight-ADRC undergo longitudinal cognitive, neuropsychologic, imaging and biomarker assessments including Clinical Dementia Rating (CDR). Among individuals with CSF and plasma data, AD cases corresponded to those with a diagnosis of dementia of the Alzheimer’s type (DAT) using criteria equivalent to the National Institute of Neurological and Communication Disorders and Stroke-Alzheimer’s Disease and Related Disorders Association for probable AD, and AD severity was determined using the Clinical Dementia Rating (CDR) at the time of lumbar puncture (for CSF samples) or blood draw (for plasma samples). Controls received the same assessment as the cases but were nondemented (CDR = 0).

Blood samples were collected in EDTA tubes (Becton Dickinson vacutainer purple top) at the visit time, immediately centrifuged at 1,500*g* for 10 min, aliquoted on two-dimensional barcoded Micronic tubes (200 ul per aliquot) and stored at −80 °C. The plasma was stored in monitored −80 °C freezer until it was pulled and sent to Somalogic (SOMAscan7k) for data generation. Proteomics data from 2,112 plasma samples from each of 2,122 participants were included in this study.

CSF samples were collected through lumbar puncture from participants after an overnight fast. Samples were processed and stored at -80 ⁰C until they were sent for protein measurement. Proteomics data from 927 CSF samples from each of 927 participants were included in this study. CSF samples from Knight-ADRC, ADNI, and DIAN were sent for proteomics using the SOMAscan platform (SOMAscan7k) in the same batch. CSF Aβ_42_, Aβ_40_, and pTau181 were measured using the LUMIPULSE *G*1200 immunoassay platform according to the manufacturer’s specifications.

The Institutional Review Board of Washington University School of Medicine in St. Louis approved the study and research was performed in accordance with the approved protocols.

#### Alzheimer’s Disease Neuroimaging Initiative (ADNI)

ADNI is a longitudinal multi-center study designed to develop early biomarkers of AD. All data used in this study were accessed from the ADNI database https://adni.loni.usc.edu/. Comprehensive details on study design, data acquisition, ethics, and policies can be found above. Proteomics data from 725 CSF samples from each of 725 participants were included in this study.

#### Dominantly Inherited Alzheimer Network (DIAN)

DIAN, led by Washington University School of Medicine in St. Louis, is a family-based long-term observational study designed to understand the earliest changes of autosomal dominant AD (ADAD). Comprehensive details on study design, data acquisition, ethics, and policies can be found at https://dian.wustl.edu/. The data used in this study are from data freeze 15 (DF15). Proteomics data from 455 CSF samples from each of 455 participants were included in this study.

#### BioFINDER2

BioFINDER2 is a Swedish prospective cohort study (<u>NCT03174938</u>) on age-related neurodegenerative diseases. Proteomics data from a total of n=848 participants, consisting of n=480 cognitively unimpaired, n=213 with mild cognitive impairment and n=155 with AD dementia were included in this study. CDR-Global scores were not measured in BioFINDER2, so for estimation participants were subdivided according to clinical diagnosis and MMSE terciles for dementia severity: cognitively normal = CDR 0, mild cognitive impairment = CDR 0.5, mild dementia (tercile 1, MMSE 22-29) = CDR 1, moderate dementia (tercile 2, MMSE 20-22) = CDR 2 and severe dementia (tercile 3, MMSE 8-19) = CDR 3. The participants were recruited at Skåne University Hospital and the Hospital of Ängelholm, Sweden. The study was approved by the Regional Ethical Committee in Lund, Sweden, and all participants gave written informed consent.

CSF samples were collected close in time after baseline clinical examination and handled according to established preanalytical protocols, previously described in detail^54^. All analyses were performed by technicians blinded to all clinical and imaging data. CSF P-tau181, Aβ42 and Aβ40 was measured using Elecsys assays in accordance with the manufacturer’s instructions (Roche Diagnostics International Ltd). CSF Aβ42/Aβ40 was used to define Aβ positivity according to previously established cutoffs of < 0.08^55^. CSF samples from the BioFINDER2 cohort were analyzed with liquid chromatography-tandem mass spectrometry (LC-MS/MS), previously described in detail^34^.

Tau-PET was performed using [^18^F]RO948. Standardized uptake value ratio (SUVR) images were created for the 70-90 min post-injection interval using the inferior cerebellar cortex as reference region. A composite corresponding to a Braak I-IV meta-region of interest was used to represent AD-related tau tangle pathology.

#### Kuopio University Hospital

The Kuopio Normal Pressure Hydrocephalus (NPH) and AD Registry and Tissue Bank includes patients from the Eastern Finnish population who were referred to the KUH neurosurgical unit for suspected NPH. The registry’s inclusive criteria encompass a wide range of hydrocephalic conditions and comorbidities: patients must exhibit one to three symptoms potentially associated with NPH (such as impaired gait, cognition, or urinary continence) along with enlarged brain ventricles (Evans’ index > 0.3) as seen on computer tomography (CT) or magnetic resonance imaging (MRI), and no other clear cause that alone explains the observed findings and symptoms. Preoperative comorbidities and conditions were recorded at baseline, and patients underwent a systematic differential diagnostic workup followed by a CSF tap test paired with gait evaluation. Follow-up was conducted on all operated patients, with optimal shunt function ensured through valve adjustment, brain imaging, shunt valve tapping, lumbar infusion testing, and shunt revision if necessary. CDR-Global scores were not measured in Kuopio, so the CERAD cognitive score was used instead.

Lumbar CSF proteomics was performed using high-throughput tandem mass tag (TMT)-labelling mass spectrometry, previously described in detail^56^. Data from 90 subjects with CSF proteomics and cognitive scoring performed within 1 year were included in this study.

The study was conducted according to the Declaration of Helsinki and all patients provided informed consent. The Research Ethics Committee of the Northern Savo Hospital District (decision No 276/13.02.00/2016).

#### Religious Order Study and Rush Memory and Aging Project (ROSMAP)

All ROSMAP participants enrolled without known dementia and agreed to detailed clinical evaluation and brain donation at death^57^. Both studies were approved by an Institutional Review Board of Rush University Medical Center (ROS IRB# L91020181, MAP IRB# L86121802). Both studies were conducted according to the principles expressed in the Declaration of Helsinki. Each participant signed an informed consent, Anatomic Gift Act, and an RADC Repository consent (IRB# L99032481) allowing their data and biospecimens to be repurposed. All participants have blood draw as a home visit, with most annual. For plasma, blood is drawn in a lavender (purple) top EDTA tube. For out of town sites, they were spun, aliquoted into nunc vials, stored in dry ice and sent to the RADC where they were transferred to -80°C. In town were brought to the RADC laboratory and processed there with the same procedures. A total of 1046 55ul samples were shipped to Stanford, then to Somalogic for proteomics (SOMAscan7k). 973 samples passed quality control.

Clinical and neuropathologic data collection has been reported in detail^10,58–61^. CDR-Global scores were not measured in ROSMAP, so for estimation participants were subdivided according to Global Cognition z-scores: cognitively normal = z-score>0, mild cognitive impairment = –1<z-score<0, mild dementia = –2<z-score<=–1, moderate dementia –3<z-score<=–2, and severe dementia z-score<=–3. These cutoffs were set based on distributions of Global Cognition z-scores per clinical diagnosis. Details on cognitive scores, neuropathology, and other patient info are described at https://www.radc.rush.edu/documentation.htm. Proteomics data from 973 plasma samples from each of 890 participants were included in this study.

### PROTEOMICS

The SomaLogic (https://somalogic.com/) SomaScan assay^62,63^, which uses slow off-rate modified DNA aptamers (SOMAmers) to bind target proteins with high specificity, was used to quantify the relative concentration of thousands of human proteins in plasma and CSF in the Stanford, Knight-ADRC, ADNI, DIAN, and ROSMAP cohorts. The v4.1 (∼7,000 proteins) assay was used for all of the mentioned cohorts and samples, except for Stanford CSF, for which the v4.0 (∼5,000 proteins) assay was used. Standard Somalogic normalization, calibration, and quality control were performed on all samples, resulting in protein measurements in relative fluorescence units (RFU). Plasma samples were further normalized to a pooled reference using an adaptive maximum likelihood procedure. The resulting values are the provided data from Somalogic and are considered “raw” data. We further performed log-10 normalization, as the assay has an expected log-normal distribution. No cohort batch corrections were applied. Acetylcholinesterase (ACHE) was removed before analyses as it confounds with AChE inhibitor treatment. CSF samples from the BioFINDER2 cohort were analyzed with liquid chromatography-tandem mass spectrometry (LC-MS/MS), previously described in detail^34^. CSF samples from the Kuopio cohort were analyzed with high-throughput tandem mass tag (TMT)-labelling mass spectrometry, previously described in detail^56^.

### COGNITIVE IMPAIRMENT STAGE CLASSIFICATION

Cognitive impairment stages reflect global clinical dementia rating (CDR-Global) scores. CDR scores of 0, 0.5, 1, 2, 3 are synonymous with cognitive impairment stages none, MCI, mild dementia, moderate dementia, and severe dementia, respectively. Stanford, Knight-ADRC, ADNI, and DIAN cohorts measured CDR-Global scores. BioFINDER2, ROSMAP, and Kuopio did not measure CDR-Global scores, so we estimated CDR-Global scores based on cognitive battery tests and clinical diagnoses as described in the sections for each cohort.

### A+T_1_+ VERSUS A–T_1_– CLASSIFICATION

Typically, “A” positivity is defined by levels of Aβ42 and “T_1_” positivity by PTau181, using a separate Gaussian Mixture model for each biomarker to derive value cutoffs^64^ (**Supplementary Fig. 1a)**. This leads to four possible groups A–T_1_–, A+T_1_–, A–T_1_+, and A+T_1_+. However, this classification system does not fit the “shape” of the data and artificially increases the number of A–T_1_+ individuals^65^ (**Supplementary Fig. 1a**), as the frequency of A–T+ individuals based on PET imaging biomarkers (the gold standards) are extremely rare^65^. To overcome this limitation, we use the CSF PTau181:Aβ42 ratio, which better fits the shape of the data (**Supplementary Fig. 1b**), to define A–T_1_– versus A+T_1_+ status (log_10_ PTau181:Aβ42 cutoff = –1; **Supplementary Fig. 1c**). Previous studies have also shown that PTau181:Aβ42 appropriately captures A–T_1_– versus A+T_1_+ status^15,16^. Aβ positivity, regardless of T status, was determined using gold-standards CSF Aβ42:Aβ40 ratio or Aβ PET.

### STATISTICAL ANALYSES

While some cohorts included multiple plasma samples from the same individual (precise numbers described in cohort sections), all analyses in this study were performed using proteomics data from only a single time point per individual. Only one CSF sample was collected per individual. For cross-sectional associations with cognitive impairment, the most recent plasma sample was used to maximize the sample size of dementia cases, which were fewer than cognitively normal cases. For analyses involving prediction of future cognitive decline from a cognitively normal-early AD baseline, the earliest plasma sample was used to maximize sample size.

#### Linear regression

The OLS function from the statsmodels^66^ Python package was used to assess linear associations between protein levels and cognitive impairment. For the unbiased proteome wide association tests in **Fig. 1b**, we tested the following linear model for each protein: cognitive impairment ∼ protein + CSF PTau181:Aβ42 + age + sex + *APOE4* dose + cohort + principal component 1 (PC1). We included PC1 of the proteome as a covariate, because previous studies have shown that it represents a large source of non-disease related variance, potentially related to heterogeneity in CSF production/clearance rates^65,67^. Inclusion of PC1 “denoised” the data greatly improved the significance of protein associations with cognitive impairment in every cohort we assessed. Multiple hypothesis testing correction was applied using the Benjamini-Hochberg method, and the significance threshold was set at a 5% false discovery rate (q-value <0.05). All other linear regression analyses in the manuscript relied on the same OLS function. Precise covariates used per analysis are displayed in the figures or described in the main text.

#### Cox proportional hazards regression

The CoxPHFitter function from the lifelines^68^ Python package was used to assess the associations between CSF YWHAG:NPTX2 and future cognitive decline (**Fig. 3e-h**, **Fig. 4i-l**). An event of cognitive decline was defined as a stage increase in cognitive impairment (i.e. none → MCI, or MCI → mild dementia). An event of conversion from cognitively normal to dementia was defined as a two stage or more increase in cognitive impairment from a cognitively normal baseline (none → mild dementia). Additional covariates included baseline age, sex, *APOE4* dose, CSF PTau181:Aβ42, and cognitive impairment, depending on the analysis. Precise covariates used per analysis are displayed in the figures or described in the text.

#### Derivation of CSF YWHAG:NPTX2 ratio

The LassoCV function from the scikit-learn^69^ Python package was used to train, in the ADNI cohort, a penalized linear model to predict cognitive impairment severity based on the levels of 214 synapse proteins that significantly changed with cognitive impairment in the ADNI and Knight-ADRC cohorts. 5-fold cross validation was implemented to identify the optimal lambda parameter. The RFECV and RFE functions from scikit-learn^69^ were used to perform recursive feature elimination on the LassoCV model to further simplify the model to facilitate clinical applications. RFECV showed that two proteins sufficiently captured the majority of the signal in the model. RFE was used to derive a model with two proteins, which resulted in the normalized ratio between YWHAG and NPTX2.

#### Derivation of plasma signature of cognitive impairment

The LassoCV function from the scikit-learn^69^ Python package was used to train, in the Knight-ADRC and ROSMAP cohorts, a penalized linear model to predict cognitive impairment severity based on the levels of 745 plasma proteins. 5-fold cross validation was implemented to identify the optimal lambda parameter. We call this model the “plasma signature” throughout the manuscript. See **Supplementary Methods** for details.

## DATA AVAILABILITY

All Stanford (including ADRC, SAMS, BPD, SCMD) data are available upon reasonable request to the Stanford-ADRC data release committee, https://web.stanford.edu/group/adrc/cgi-bin/web-proj/datareq.php. Data from specific cohorts can be requested to the following cohort leaders: ADRC, Tony Wyss-Coray; SAMS, Beth Mormino or Anthony Wagner; BPD and SCMD, Kathleen Poston. Knight-ADRC proteomics data were generated by the laboratory of principal investigator Carlos Cruchaga and are available upon reasonable request to The National Institute on Aging Genetics of Alzheimer’s Disease Data Storage Site (NIAGADS) https://www.niagads.org/knight-adrc-collection. ADNI data can be requested at the ADNI database (https://adni.loni.usc.edu/). DIAN data can be requested at https://dian.wustl.edu/our-research/for-investigators/diantu-investigator-resources/dian-tu-biospecimen-request-form/. Pseudonymized BioFINDER2 data can be shared to qualified academic researchers after request to cohort leader Oskar Hansson for the purpose of replicating procedures and results presented in the study. Data transfer must be performed in agreement with EU legislation regarding general data protection regulation and decisions by the Ethical Review Board of Sweden and Region Skåne. Kuopio data can be requested and accessed via a repository on Terra https://app.terra.bio/#workspaces/marsh-terra-inph/iNPH_Proteomics_Workspace. ROSMAP data can be requested at https://www.radc.rush.edu and www.synpase.org.

## CODE AVAILABILITY

The CSF YWHAG.1:NPTX2 ratio can be derived by log_10_ normalizing protein levels (YWHAG.1 SeqId=4179-57, NPTX2 SeqId=6521-35), z-score normalizing using means and standard deviations from our cohorts (YWHAG.1 mean=3.425, std=0.183; NPTX2 mean=4.099, std=0.171), then taking the difference between normalized YWHAG.1 and NPTX2 values.

The plasma signature of cognitive impairment is a linear model (linear combination of protein values and weights with final addition of an intercept value) can be accessed in a Python package called “plasmaCI” (at the time of publication) to easily derive plasma signature values from any SomaScan (assay v4 and above) plasma proteomics sample. Two separate plasma signatures were trained, one using a pre-defined set of 745 proteins (See **Supplementary Methods**) on the v4.1 assay (∼7,000 proteins) and another using 552 of the 745 pre-defined proteins detected on the v4 assay (∼5,000 proteins). The v4 and v4.1 signatures are highly correlated (r=0.97).

Protein weights for both versions of the plasma signature are provided in **Supplementary Table 7**. Before applying model weights, plasma protein levels should be log_10_ normalized, then z-score normalized using means and standard deviations from our training data (**Supplementary Table 7**). Additionally, if using the v4 assay and signature, simple multiplicative scale factors should be applied before log_10_ and z-score normalization (**Supplementary Table 7**).

## AUTHOR CONTRIBUTIONS

H.S.O. conceptualized the study. H.S.O. led study design and analyses. D.Y.U. aided in study design and analyses. L.K. aided in analyses in the BioFINDER2 cohort. Z.Z. aided in analyses in the ADNI cohort. Y.S. aided in analyses in the DIAN cohort. A.F. aided in analyses in the Stanford-ADRC cohort. N.M. aided in analyses in the Kuopio cohort. J.T., I.G., C.Y., D.W., M.A., Y.L.G., and A.T. provided data, aided in analyses, and/or provided insights. T.R., S.-K. H., M.H., A.Li., and A.Lu., collected data from the Kuopio cohort. K.P. established the Stanford-BPD cohort. E.M. and A.D.W. established the SAMS cohort. E.N.W. led fluid AD biomarker data collection in Stanford cohorts. D.C. led plasma collection in the Stanford cohorts. V.L., B.S., and H.Z., established the Kuopio cohort. A.J.E. provided key insights on the Alzheimer’s field. D.B. established the ROSMAP cohort. N.F. provided key insights on the Alzheimer’s field and aided in analyses the ADNI cohort. O.H. established the BioFINDER2 cohort. C.C. established the Knight-ADRC cohort. T.W.-C. established the Stanford-ADRC cohort. K.P., E.M., A.D.W., E.N.W., V.L., B.S., H.Z., D.B., N.F., O.H., C.C., provided data and insights. H.S.O. produced figures and wrote the manuscript. T.W.-C. edited the manuscript. H.S.O. and T.W.-C supervised the study. All authors critically revised the manuscript for intellectual content. All authors read and approved the final version of the manuscript.

## ACKNOWLEDGEMENTS

We thank B. Lehallier, J. Rutledge, L. Gold, and members of the Wyss-Coray laboratory for feedback and support and D. Channappa for laboratory management. We are also grateful for the help of Marita Parviainen and Tiina Laaksonen with patient management and cognitive testing.

This work was supported by the Stanford Alzheimer’s Disease Research Center (National Institute on Aging grants P50AG047366 and P30AG066515), the National Institute on Aging (AG072255, T.W.-C), the Milky Way Research Foundation (T.W.-C.), the Knight Initiative for Brain Resilience (T.W.-C.), the Stanford Graduate Fellowship (H.S.O.), and the National Science Foundation Graduate Research Fellowship (H.S.O.). E.N.W. is supported by a grant from the KIBR. Y.L.G is supported by the Stanford’s Center for Clinical and Translational Education and Research award, under the Biostatistics, Epidemiology and Research Design (BERD) Program: UL1TR003142.

This work was also supported by grants from the National Institutes of Health (R01AG044546 (CC), P01AG003991(CC, JCM), RF1AG053303 (CC), RF1AG058501 (CC), U01AG058922 (CC), RF1AG074007 (YJS)), the Chan Zuckerberg Initiative (CZI), the Michael J. Fox Foundation (LI, CC), the Department of Defense (LI-W81XWH2010849), the Alzheimer’s Association Zenith Fellows Award (ZEN-22-848604, awarded to CC), and an Anonymous foundation. The recruitment and clinical characterization of research participants at Washington University were supported by NIH P30AG066444 (JCM), P01AG03991(JCM), and P01AG026276(JCM). This work was supported by access to equipment made possible by the Hope Center for Neurological Disorders, the Neurogenomics and Informatics Center (NGI: https://neurogenomics.wustl.edu/) and the Departments of Neurology and Psychiatry at Washington University School of Medicine.

The BioFINDER-2 study was supported by European Research Council (ADG-101096455), Alzheimer’s Association (ZEN24-1069572, SG-23-1061717), GHR Foundation, Swedish Research Council (2022-00775), ERA PerMed (ERAPERMED2021-184), Knut and Alice Wallenberg foundation (2022-0231), Strategic Research Area MultiPark (Multidisciplinary Research in Parkinson’s disease) at Lund University, Swedish Alzheimer Foundation (AF-980907), Swedish Brain Foundation (FO2021-0293), Parkinson foundation of Sweden (1412/22), Cure Alzheimer’s fund, Rönström Family Foundation, Konung Gustaf V:s och Drottning Victorias Frimurarestiftelse, Skåne University Hospital Foundation (2020-O000028), Regionalt Forskningsstöd (2022-1259) and Swedish federal government under the ALF agreement (2022-Projekt0080).

SAMS is supported by grants from the National Institutes of Health (R01AG048076, R21AG058859), Stanford Wu Tsai Neurosciences Institute, and Stanford Center for Precision Health and Integrated Diagnostics (PHIND).

ROSMAP is supported by P30AG10161, P30AG72975, R01AG15819, R01AG17917, U01AG46152, and U01AG61356.

The Kuopio study was funded by the Alzheimer’s Association, Academy of Finland (grant numbers 338182, 328287), KUH VTR Fund, Sigrid Juselius Foundation, the Strategic Neuroscience Funding of the University of Eastern Finland, and Alzheimer’s Association ADSF-24-1284326-C.

H.Z. is a Wallenberg Scholar and a Distinguished Professor at the Swedish Research Council supported by grants from the Swedish Research Council (#2023-00356; #2022-01018 and #2019-02397), the European Union’s Horizon Europe research and innovation programme under grant agreement No 101053962, Swedish State Support for Clinical Research (#ALFGBG-71320), the Alzheimer Drug Discovery Foundation (ADDF), USA (#201809-2016862), the AD Strategic Fund and the Alzheimer’s Association (#ADSF-21-831376-C, #ADSF-21-831381-C, #ADSF-21-831377-C, and #ADSF-24-1284328-C), the European Partnership on Metrology, co-financed from the European Union’s Horizon Europe Research and Innovation Programme and by the Participating States (NEuroBioStand, #22HLT07), the Bluefield Project, Cure Alzheimer’s Fund, the Olav Thon Foundation, the Erling-Persson Family Foundation, Familjen Rönströms Stiftelse, Stiftelsen för Gamla Tjänarinnor, Hjärnfonden, Sweden (#FO2022-0270), the European Union’s Horizon 2020 research and innovation programme under the Marie Skłodowska-Curie grant agreement No 860197 (MIRIADE), the European Union Joint Programme – Neurodegenerative Disease Research (JPND2021-00694), the National Institute for Health and Care Research University College London Hospitals Biomedical Research Centre, and the UK Dementia Research Institute at UCL (UKDRI-1003).

B.S. was supported by the Alzheimer’s Association (ADSF-21-836089-C, ADSF-21-836083-C, ADSF-21-836085-C). N.M. was supported by NIH training grants 5T32AG222-30 and 1F32AG079666-01.

## EXTENDED DATA FIGURES

**Extended Data Figure 1.**
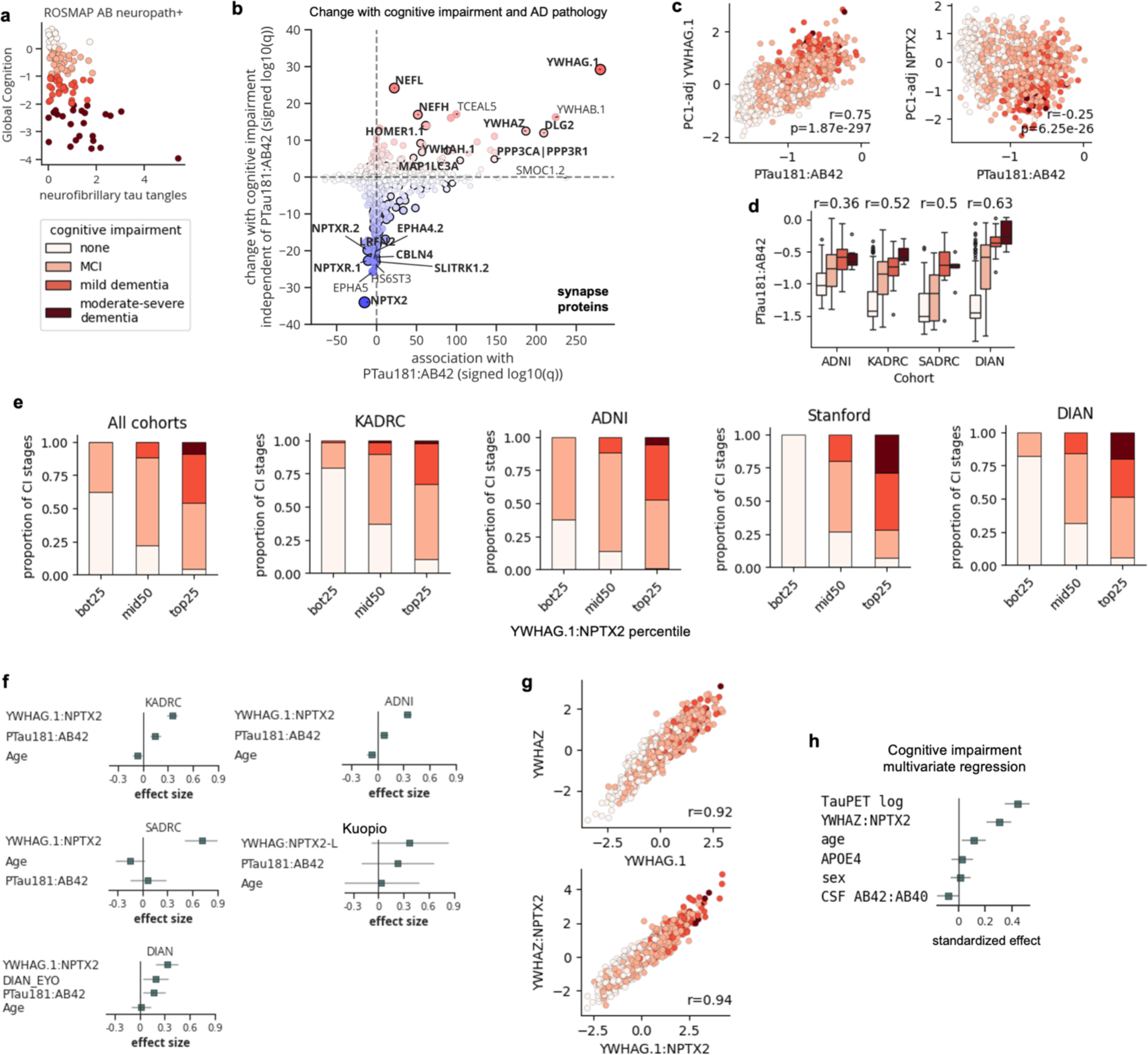
CSF YWHAG:NPTX2 ratio explains a substantial proportion of variance in cognitive impairment beyond amyloid and tau in AD. **a,** Global cognition score versus tau tangle load in Aβ+ individuals in the ROSMAP cohort. Aβ and tau do not sufficiently explain cognitive impairment. **b,** Scatterplot showing both change with cognitive impairment independent of PTau181:Aβ42 (y-axis) and association with PTau181:Aβ42 (x-axis). Axes show signed –log10 q-values (Benjamini-Hochberg adjusted p-value). Bold indicates synapse proteins. **c,** Scatterplot showing PC1-adjusted YWHAG.1 (left) and NPTX2 (right) versus PTau181:Aβ42, colored by cognitive impairment. **d,** Boxplot showing PTau181:Aβ42 versus cognitive impairment severity across cohorts. Standard boxplot metrics used. **e,** Stacked bar plot showing proportions of different cognitive impairment stages among different YWHAG:NPTX2 percentile groups, in all and each cohort. **f,** Results from linear models regressing cognitive impairment against the displayed covariates, per cohort. Standardizes effects and 95% confidence intervals shown. **g,** Scatterplot showing YWHAZ versus YWHAG.1, colored by cognitive impairment (left). Scatterplot showing YWHAZ:NPTX2 versus YWHAG.1:NPTX2, colored by cognitive impairment (right). **j,** Results from a multivariate linear model regressing cognitive impairment against the displayed covariates in the BioFINDER2 cohort. Standardized betas and 95% confidence intervals shown.

**Extended Data Figure 2.**
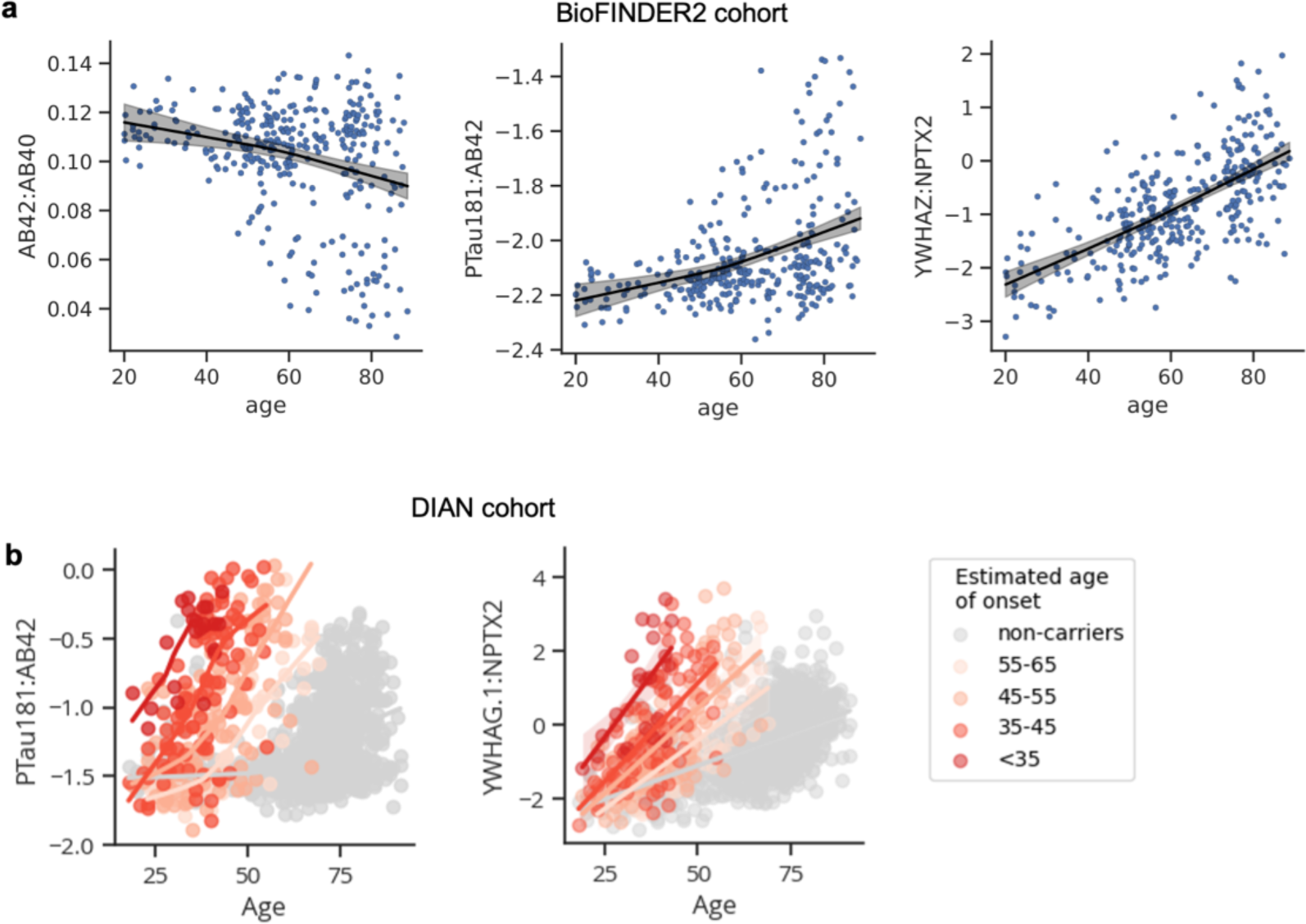
Changes in CSF YWHAG:NPTX2 with normal aging and ADAD. **a,** Scatterplots showing changes in CSF Aβ42:Aβ40, PTau181:Aβ42, and YWHAG.1:NPTX2 with cognitively normal aging in the BioFINDER2 cohort. **a,** Scatterplots showing changes in CSF PTau181:Aβ42, and YWHAG.1:NPTX2 with age in ADAD mutation carriers, binned by estimated age of onset, in the DIAN cohort.

## REFERENCES

1. Knopman, D. S. et al. Alzheimer disease. Nat. Rev. Dis. Primer 7, 33 (2021).

2. Dujardin, S. et al. Tau molecular diversity contributes to clinical heterogeneity in Alzheimer’s disease. Nat. Med. 26, 1256–1263 (2020).

3. Ossenkoppele, R. et al. Amyloid and tau PET-positive cognitively unimpaired individuals are at high risk for future cognitive decline. Nat. Med. 28, 2381–2387 (2022).

4. Strikwerda-Brown, C. et al. Association of Elevated Amyloid and Tau Positron Emission Tomography Signal With Near-Term Development of Alzheimer Disease Symptoms in Older Adults Without Cognitive Impairment. JAMA Neurol. 79, 975–985 (2022).

5. Zetterberg, H. & Bendlin, B. B. Biomarkers for Alzheimer’s disease—preparing for a new era of disease-modifying therapies. Mol. Psychiatry 26, 296–308 (2021).

6. van Dyck Christopher H., et al. Lecanemab in Early Alzheimer’s Disease. N. Engl. J. Med. 388, 9–21 (2023).

7. Hansson, O. Biomarkers for neurodegenerative diseases. Nat. Med. 27, 954–963 (2021).

8. Mostafavi, S. et al. A molecular network of the aging human brain provides insights into the pathology and cognitive decline of Alzheimer’s disease. Nat. Neurosci. 21, 811–819 (2018).

9. Hanseeuw, B. J. et al. Association of Amyloid and Tau With Cognition in Preclinical Alzheimer Disease: A Longitudinal Study. JAMA Neurol. 76, 915–924 (2019).

10. Boyle, P. A. et al. To what degree is late life cognitive decline driven by age-related neuropathologies? Brain 144, 2166–2175 (2021).

11. Tosun, D. et al. Contribution of Alzheimer’s biomarkers and risk factors to cognitive impairment and decline across the Alzheimer’s disease continuum. Alzheimers Dement. 18, 1370–1382 (2022).

12. Andersen, S. L. Centenarians as Models of Resistance and Resilience to Alzheimer’s Disease and Related Dementias. Adv. Geriatr. Med. Res. 2, e200018 (2020).

13. Zhang, M. et al. The correlation between neuropathology levels and cognitive performance in centenarians. Alzheimers Dement. 19, 5036–5047 (2023).

14. Jack Jr., C. R., et al. Revised criteria for diagnosis and staging of Alzheimer’s disease: Alzheimer’s Association Workgroup. Alzheimers Dement. n/a, (2024).

15. Martínez-Dubarbie, F. et al. Accuracy of plasma Aβ40, Aβ42, and p-tau181 to detect CSF Alzheimer’s pathological changes in cognitively unimpaired subjects using the Lumipulse automated platform. Alzheimers Res. Ther. 15, 163 (2023).

16. Motta, C. et al. Different associations between amyloid-βeta 42, amyloid-βeta 40, and amyloid-βeta 42/40 with soluble phosphorylated-tau and disease burden in Alzheimer’s disease: a cerebrospinal fluid and fluorodeoxyglucose-positron emission tomography study. Alzheimers Res. Ther. 15, 144 (2023).

17. Horie, K. et al. CSF MTBR-tau243 is a specific biomarker of tau tangle pathology in Alzheimer’s disease. Nat. Med. 29, 1954–1963 (2023).

18. van der Flier, W. M. & Scheltens, P. The ATN Framework—Moving Preclinical Alzheimer Disease to Clinical Relevance. JAMA Neurol. 79, 968–970 (2022).

19. Koopmans, F. et al. SynGO: An Evidence-Based, Expert-Curated Knowledge Base for the Synapse. Neuron 103, 217–234.e4 (2019).

20. Zhang, J. & Zhou, Y. 14-3-3 Proteins in Glutamatergic Synapses. Neural Plast. 2018, 8407609 (2018).

21. Segal, D. et al. A central chaperone-like role for 14-3-3 proteins in human cells. Mol. Cell 83, 974–993.e15 (2023).

22. Ali, M. & et. al. Multi-cohort cerebrospinal fluid proteomics identifies robust molecular signatures for asymptomatic and symptomatic Alzheimer’s disease,. (2024).

23. Stallings, N. R. et al. Pin1 mediates Aβ42-induced dendritic spine loss. Sci. Signal. 11, eaap8734 (2018).

24. Yin, Y., et al. Tau accumulation induces synaptic impairment and memory deficit by calcineurin-mediated inactivation of nuclear CaMKIV/CREB signaling. Proc. Natl. Acad. Sci. 113, E3773–E3781 (2016).

25. Stallings, N. R., O’Neal, M. A., Hu, J., Shen, Z.-J. & Malter, J. S. Long-term normalization of calcineurin activity in model mice rescues Pin1 and attenuates Alzheimer’s phenotypes without blocking peripheral T cell IL-2 response. Alzheimers Res. Ther. 15, 179 (2023).

26. Drummond, E. et al. The amyloid plaque proteome in early onset Alzheimer’s disease and Down syndrome. Acta Neuropathol. Commun. 10, 53 (2022).

27. Johnson, E. C. B. et al. Cerebrospinal fluid proteomics define the natural history of autosomal dominant Alzheimer’s disease. Nat. Med. 29, 1979–1988 (2023).

28. Chapman, G., Shanmugalingam, U. & Smith, P. D. The Role of Neuronal Pentraxin 2 (NP2) in Regulating Glutamatergic Signaling and Neuropathology. Front. Cell. Neurosci. 13, (2020).

29. Pelkey, K. A. et al. Pentraxins Coordinate Excitatory Synapse Maturation and Circuit Integration of Parvalbumin Interneurons. Neuron 85, 1257–1272 (2015).

30. Gabitto, M., Travaglini, K., Jeannelle, A. & et. al. Integrated multimodal cell atlas of Alzheimer’s disease. (2023).

31. Mathys, H. et al. Single-cell atlas reveals correlates of high cognitive function, dementia, and resilience to Alzheimer’s disease pathology. Cell 186, 4365–4385.e27 (2023).

32. Zhou, J. et al. The neuronal pentraxin Nptx2 regulates complement activity and restrains microglia-mediated synapse loss in neurodegeneration. Sci. Transl. Med. 15, eadf0141 (2023).

33. Sathe, G. et al. Quantitative Proteomic Profiling of Cerebrospinal Fluid to Identify Candidate Biomarkers for Alzheimer’s Disease. PROTEOMICS – Clin. Appl. 13, 1800105 (2019).

34. Nilsson, J. et al. Cerebrospinal fluid biomarker panel for synaptic dysfunction in a broad spectrum of neurodegenerative diseases. Brain awae032 (2024) doi:10.1093/brain/awae032.

35. Jiang, Y. et al. Large-scale plasma proteomic profiling identifies a high-performance biomarker panel for Alzheimer’s disease screening and staging. Alzheimers Dement. 18, 88– 102 (2022).

36. Ryman, D. C. et al. Symptom onset in autosomal dominant Alzheimer disease. Neurology 83, 253–260 (2014).

37. Mitchell, A. J. & Shiri-Feshki, M. Rate of progression of mild cognitive impairment to dementia – meta-analysis of 41 robust inception cohort studies. Acta Psychiatr. Scand. 119, 252–265 (2009).

38. Jia Jianping et al. Biomarker Changes during 20 Years Preceding Alzheimer’s Disease. N. Engl. J. Med. 390, 712–722 (2024).

39. Pooler, A. M. et al. Amyloid accelerates tau propagation and toxicity in a model of early Alzheimer’s disease. Acta Neuropathol. Commun. 3, 14 (2015).

40. Wu, J. W. et al. Neuronal activity enhances tau propagation and tau pathology in vivo. Nat. Neurosci. 19, 1085–1092 (2016).

41. Franzmeier, N. et al. Elevated CSF GAP-43 is associated with accelerated tau accumulation and spread in Alzheimer’s disease. Nat. Commun. 15, 202 (2024).

42. Gómez de San José, N., et al. Neuronal pentraxins as biomarkers of synaptic activity: from physiological functions to pathological changes in neurodegeneration. J. Neural Transm. 129, 207–230 (2022).

43. Nilsson, J. et al. Cerebrospinal fluid biomarker panel for synaptic dysfunction in Alzheimer’s disease. Alzheimers Dement. Diagn. Assess. Dis. Monit. 13, e12179 (2021).

44. Ye, X.-G. et al. YWHAG Mutations Cause Childhood Myoclonic Epilepsy and Febrile Seizures: Molecular Sub-regional Effect and Mechanism. Front. Genet. 12, (2021).

45. Hashiguchi, M., Sobue, K. & Paudel, H. K. 14-3-3ζ Is an Effector of Tau Protein Phosphorylation*. J. Biol. Chem. 275, 25247–25254 (2000).

46. Scott-Hewitt, N., et al. Local externalization of phosphatidylserine mediates developmental synaptic pruning by microglia. EMBO J. 39, e105380 (2020).

47. Colom-Cadena, M. et al. The clinical promise of biomarkers of synapse damage or loss in Alzheimer’s disease. Alzheimers Res. Ther. 12, 21 (2020).

48. Morrison, J. H. & Baxter, M. G. The ageing cortical synapse: hallmarks and implications for cognitive decline. Nat. Rev. Neurosci. 13, 240–250 (2012).

49. Oh, H. S.-H. et al. Organ aging signatures in the plasma proteome track health and disease. Nature 624, 164–172 (2023).

50. Wilson, E. N. et al. Performance of a fully-automated Lumipulse plasma phospho-tau181 assay for Alzheimer’s disease. Alzheimers Res. Ther. 14, 172 (2022).

51. Wilson, E. N. et al. Soluble TREM2 is elevated in Parkinson’s disease subgroups with increased CSF tau. Brain 143, 932–943 (2020).

52. Plastini, M. J. et al. Multiple biomarkers improve diagnostic accuracy across Lewy body and Alzheimer’s disease spectra. Ann. Clin. Transl. Neurol. 11, 1197–1210 (2024).

53. McKhann, G. et al. Clinical diagnosis of Alzheimer’s disease. Neurology 34, 939–939 (1984).

54. Palmqvist, S. et al. Discriminative Accuracy of Plasma Phospho-tau217 for Alzheimer Disease vs Other Neurodegenerative Disorders. JAMA 324, 772–781 (2020).

55. Pichet Binette, A., et al. Amyloid-associated increases in soluble tau relate to tau aggregation rates and cognitive decline in early Alzheimer’s disease. Nat. Commun. 13, 6635 (2022).

56. Weiner, S. et al. Optimized sample preparation and data analysis for TMT proteomic analysis of cerebrospinal fluid applied to the identification of Alzheimer’s disease biomarkers. Clin. Proteomics 19, 13 (2022).

57. Bennett, D. A. et al. Religious Orders Study and Rush Memory and Aging Project. J. Alzheimers Dis. 64, S161–S189 (2018).

58. Bennett, D. A. et al. Neuropathology of older persons without cognitive impairment from two community-based studies. Neurology 66, 1837–1844 (2006).

59. Bennett, D. A. et al. Decision Rules Guiding the Clinical Diagnosis of Alzheimer’s Disease in Two Community-Based Cohort Studies Compared to Standard Practice in a Clinic-Based Cohort Study. Neuroepidemiology 27, 169–176 (2006).

60. Bennett, D. A. et al. Natural history of mild cognitive impairment in older persons. Neurology 59, 198–205 (2002).

61. Boyle, P. A. et al. Attributable risk of Alzheimer’s dementia attributed to age-related neuropathologies. Ann. Neurol. 85, 114–124 (2019).

62. Williams, S. A. et al. Plasma protein patterns as comprehensive indicators of health. Nat. Med. 25, 1851–1857 (2019).

63. Gold, L. et al. Aptamer-Based Multiplexed Proteomic Technology for Biomarker Discovery. PLOS ONE 5, e15004 (2010).

64. Timsina, J. et al. Harmonization of CSF and imaging biomarkers in Alzheimer’s disease: Need and practical applications for genetics studies and preclinical classification. Neurobiol. Dis. 190, 106373 (2024).

65. Karlsson, L. et al. Cerebrospinal fluid reference proteins increase accuracy and interpretability of biomarkers for brain diseases. Nat. Commun. 15, 3676 (2024).

66. Seabold, S. & Perktold, J. Statsmodels: Econometric and Statistical Modeling with Python. in *Proceedings of the 9th Python in Science Conference* (eds. Walt, S. van der & Millman, J.) 92–96 (2010). doi:10.25080/Majora-92bf1922-011.

67. Shen, Y. et al. Systematic proteomics in Autosomal dominant Alzheimer’s disease reveals decades-early changes of CSF proteins in neuronal death, and immune pathways. medRxiv 2024.01.12.24301242 (2024) doi:10.1101/2024.01.12.24301242.

68. Davidson-Pilon, Cameron. (2022). lifelines, survival analysis in Python (v0.27.0). Zenodo. 10.5281/zenodo.6359609.

69. Pedregosa, F. et al. Scikit-learn: Machine Learning in Python. J. Mach. Learn. Res. 12, 2825–2830 (2011).

