## Supplementary Methods for "Synapse protein signatures in cerebrospinal fluid and plasma predict cognitive maintenance versus decline in Alzheimer’s disease"

**A+T_1_+ VERSUS A–T_1_– CLASSIFICATION**

Typically, “A” positivity is defined by levels of Aβ42 and “T_1_” positivity by PTau181, using a separate Gaussian Mixture model for each biomarker to derive value cutoffs^1^ (**Supplementary Fig. 1a)**. This leads to four possible groups A–T_1_–, A+T_1_–, A–T_1_+, and A+T_1_+. However, this classification system does not fit the “shape” of the data and artificially increases the number of A–T_1_+ individuals^2^ (**Supplementary Fig. 1a**), as the frequency of A–T+ individuals based on PET imaging biomarkers (the gold standards) are extremely rare^2^. To overcome this limitation, we use the CSF PTau181:Aβ42 ratio, which better fits the shape of the data (**Supplementary Fig. 1b**), to define A–T_1_– versus A+T_1_+ status (log_10_ PTau181:Aβ42 cutoff = –1; **Supplementary Fig. 1c**). Previous studies have also shown that PTau181:Aβ42 appropriately captures A–T_1_– versus A+T_1_+ status^3,4^. Aβ positivity, regardless of T status, was determined using gold-standards CSF Aβ42:Aβ40 ratio or Aβ PET.

**Derivation of plasma signature of cognitive impairment (CI).**

We first sought to determine whether plasma YWHAG:NPTX2 could faithfully reflect CSF YWHAG:NPTX2. We used the set of 1,194 paired CSF and plasma samples (n=597 each) that were collected within 6 months of each other from the Knight-ADRC and Stanford cohorts. We found no significant correlation (**Supplementary Fig. 2a**). We then tested a series of frameworks to develop a plasma signature that could more accurately reflect CSF YWHAG:NPTX2 and CI.

First, we aimed to develop a plasma signature of CSF YWHAG:NPTX2 using the 240 plasma synapse proteins that changed with CI in the CSF, with the idea that perhaps many more synapse proteins beyond YWHAG and NPTX2 are needed to train a robust model from plasma. We split the 597 matched CSF and plasma samples into 478 for training (80%) and 119 for testing (20%). We used the LassoCV function from the scikit-learn^5^ Python package to train a penalized linear model to predict CI severity based on the levels of the 240 significant synapse proteins. Though the correlation between this plasma signature and CSF YWHAG:NPTX2 was decent, the correlations with CI were mild (**Supplementary Fig. 2b**).

To improve the robustness of the signature, we sought to use the same set of proteins to train a predictor of CI instead of CSF YWHAG:NPTX2 directly. This allowed us to dramatically increase training sample size as CI, but not matched CSF, was measured across many more samples and cohorts. We used Knight-ADRC and ROSMAP for training and Stanford for testing. This improved correlations with CI but weakened correlations with CSF YWHAG:NPTX2 (**Supplementary Fig. 2c**).

We believed there was additional potential for improvement. We sought to expand upon our modeling framework to include non-synapse plasma proteins, while also filtering out proteins that were unstable across cohorts and were likely affected by ApoE-binding (**Supplementary Fig. 3a)**. To elaborate, we first applied the Weighted Gene Co-expression Network Analysis^6^ (WGCNA) package in R to derive 14 correlation modules from the plasma proteome. Knight-ADRC and ROSMAP cohorts were used for module training and Stanford for testing. We used the “blockwiseModules” function with the following parameters: power = 5, networkType = "unsigned", corType="bicor", maxBlockSize=10000, deepSplit = 2, pamRespectsDendro = F, minModuleSize = 20, reassignThreshold=1e-6, mergeCutHeight = 0.25. Proteins not assigned to any module were assigned to the “grey” module (M0). We confirmed that modules were robust within and across cohorts, as proteins within a module tended to correlate with their respective module eigenproteins more than other module eigenproteins, and this was very consistent across train and test data (**Supplementary Fig. 3b**).

We next regressed each module eigenprotein with age, sex, cohort, *APOE4* dose, and CI (**Supplementary Fig. 4a**). We identified several modules with strong cohort-effects (**Supplementary Fig. 4b**), suggesting proteins in these modules are likely sensitive to differences in blood processing between cohorts/centers. We filtered out proteins in these modules. We also identified a single module with a strong *APOE4* effect (**Supplementary Fig. 4c**). The strong “block-like” association between *APOE* genotype and proteins in this module suggest that *APOE* genotype may alter the binding affinity between these proteins and their respective Somalogic aptamer probes, potentially due to apoE-protein interactions. Notably, NfL is a member of this module and has a strong negative *APOE4* effect, decreasing in signal with increasing *APOE4* dose. This aligns with previous reports of discordance between SomaScan and antibody-based assays for NfL^7^. We filtered out proteins in this *APOE* module. Lastly, for each of the remaining seven modules and the “grey” module (M0), we performed a Fisher’s exact test to identify modules that were enriched for synapse proteins that changed with CI independent of Aβ and tau in the CSF (**Fig. 1b**). Module 3 was the only module that was strongly enriched for these synapse proteins (**Supplementary Fig. 4d**), so we trained our plasma signature of CI using the 745 proteins in module 3.

The LassoCV function from the scikit-learn^5^ Python package was used to train a penalized linear model to predict cognitive impairment severity based on the levels of the 745 plasma proteins in module 3. The same train/test split as for WGCNA was used, with Knight-ADRC and ROSMAP cohorts for training and Stanford for testing. 5-fold cross validation was implemented to identify the optimal lambda parameter. We call this model the “plasma signature” throughout the manuscript (**Fig. 4**).

**REFERENCES**

1.	Timsina, J. *et al.* Harmonization of CSF and imaging biomarkers in Alzheimer’s disease: Need and practical applications for genetics studies and preclinical classification. *Neurobiol. Dis.* **190**, 106373 (2024).

2.	Karlsson, L. *et al.* Cerebrospinal fluid reference proteins increase accuracy and interpretability of biomarkers for brain diseases. *Nat. Commun.* **15**, 3676 (2024).

3.	Martínez-Dubarbie, F. *et al.* Accuracy of plasma Aβ40, Aβ42, and p-tau181 to detect CSF Alzheimer’s pathological changes in cognitively unimpaired subjects using the Lumipulse automated platform. *Alzheimers Res. Ther.* **15**, 163 (2023).

4.	Motta, C. *et al.* Different associations between amyloid-βeta 42, amyloid-βeta 40, and amyloid-βeta 42/40 with soluble phosphorylated-tau and disease burden in Alzheimer’s disease: a cerebrospinal fluid and fluorodeoxyglucose-positron emission tomography study. *Alzheimers Res. Ther.* **15**, 144 (2023).

5.	Pedregosa, F. *et al.* Scikit-learn: Machine Learning in Python. *J. Mach. Learn. Res.* **12**, 2825–2830 (2011).

6.	Langfelder, P. & Horvath, S. WGCNA: an R package for weighted correlation network analysis. *BMC Bioinformatics* **9**, 559 (2008).

7.	Eldjarn, G. H. *et al.* Large-scale plasma proteomics comparisons through genetics and disease associations. *Nature* **622**, 348–358 (2023).

**SUPPLEMENTARY FIGURES**

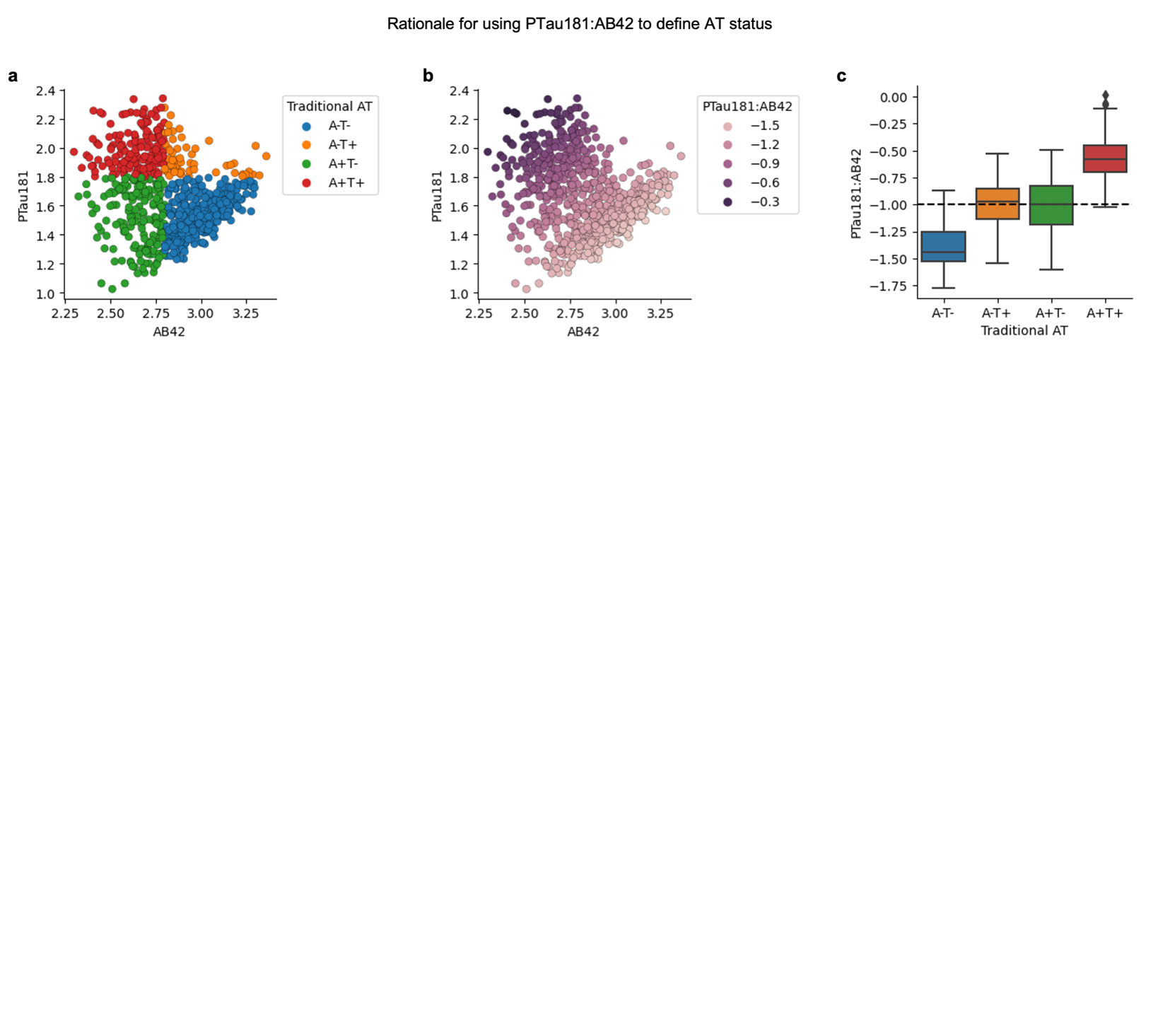

**Supplementary Figure 1. AT_1_ status classification using the CSF PTau181:Aβ42 ratio.**

**a,** Scatterplot showing CSF PTau181 versus Aβ42, colored by AT_1_ status based on traditional classification methods, which use Gaussian Mixture models to derive binary cutoffs per biomarker.

**b,** Scatterplot showing CSF PTau181 versus Aβ42, colored by CSF PTau181:Aβ42 ratio.

**c,** Boxplot showing PTau181:Aβ42 versus traditional AT_1_ status. For this study, CSF PTau181:Aβ42 > –1 was used to define A+T1+ individuals, and CSF PTau181:Aβ42 < –1 for A–T1– individuals.

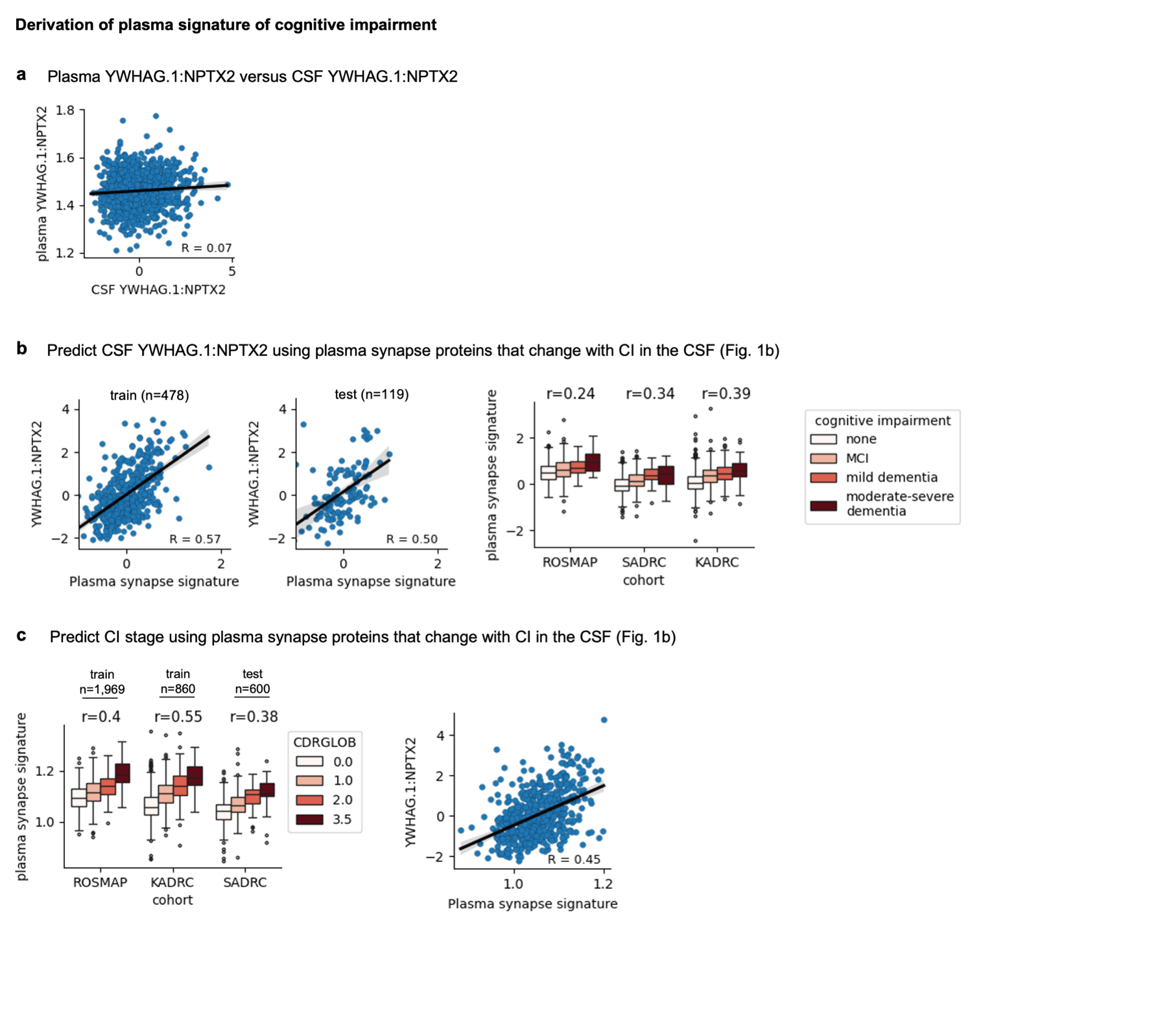
**Supplementary Figure 2. WGCNA of the plasma proteome across cohorts.**

**a,** Plasma versus CSF YWHAG:NPTX2 in samples collected within 6 months in the Knight-ADRC and Stanford cohorts.

**b,** A linear model was trained to predict CSF YWHAG:NPTX2 using plasma synapse proteins that changed with CI in the CSF (Fig. 1b). Correlations between actual (y-axis) and predicted (x-axis) CSF YWHAG:NPTX2 values are shown (left). Correlations between predicted CSF YWHAG:NPTX2 values and CI stage across cohorts are shown (right).

**c,** A linear model was trained to predict CI stage using plasma synapse proteins that changed with CI in the CSF (Fig. 1b). Correlations between actual (x-axis) and predicted (y-axis) CI stages are shown (left). Correlation between CSF YWHAG:NPTX2 and predicted CI values is shown (right).

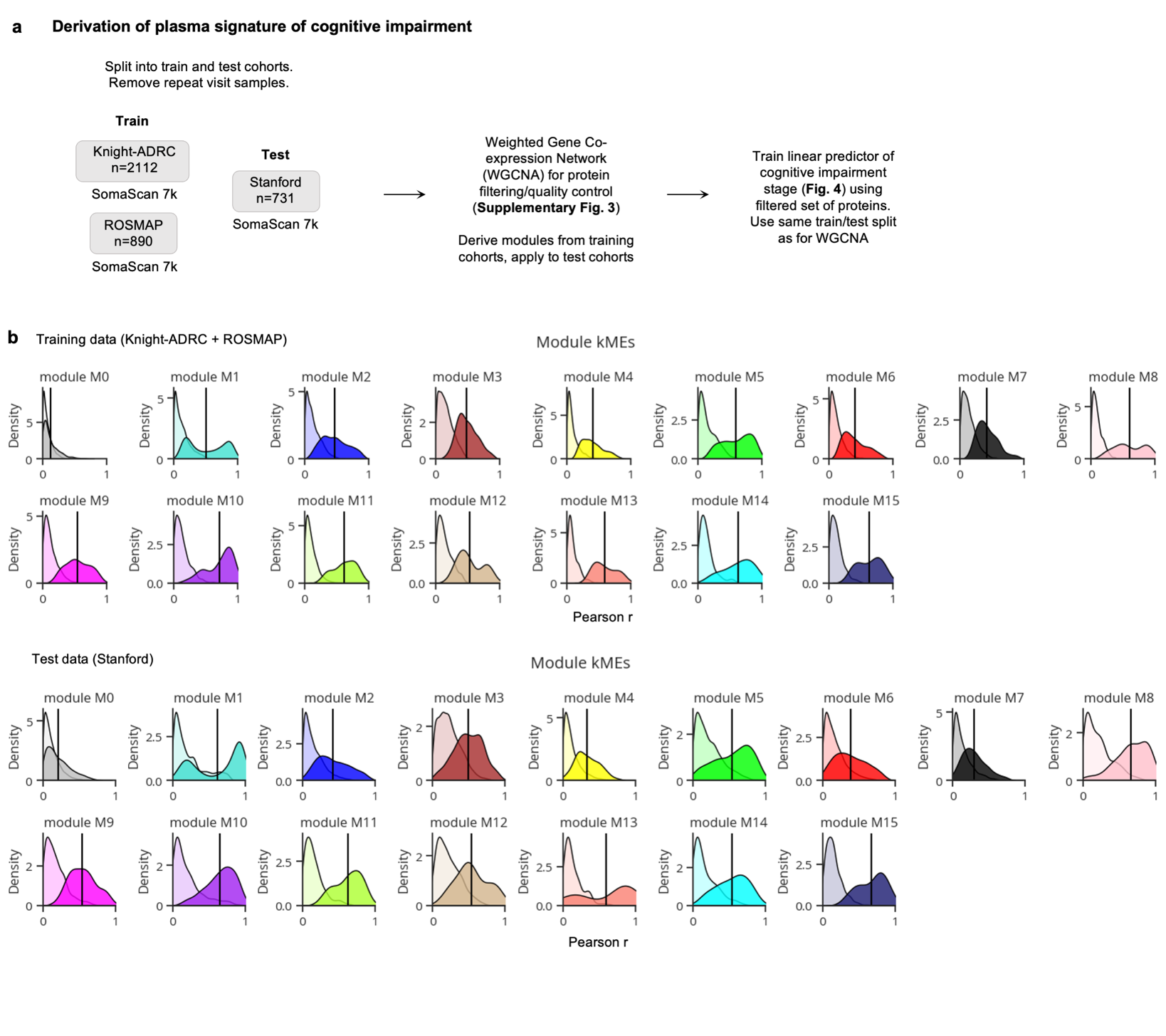

**Supplementary Figure 3. WGCNA of the plasma proteome across cohorts.**

**a,** Workflow to derive robust plasma signature of cognitive impairment.

**b,** WGCNA resulted in 14 modules. Proteins not assigned to any modules were assigned to module zero (M0). Density plots showing the distribution of correlations between proteins and module eigenproteins are shown. Correlations between proteins and their respective module eigenproteins are included in the opaque distributions, while correlations between proteins and non-matching module eigenproteins are included in the transparent distributions. Correlations from training data are shown above and testing data below.

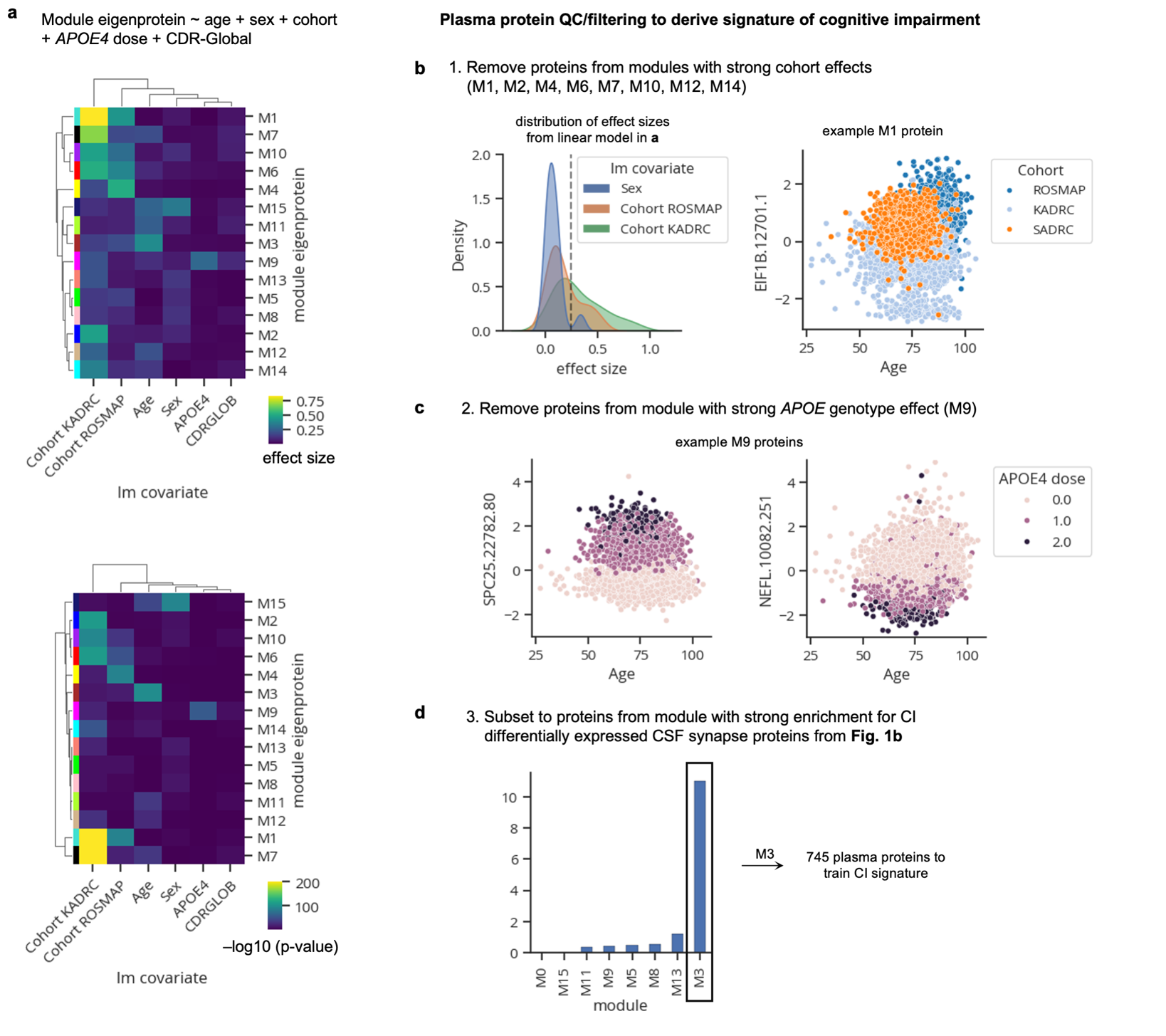

**Supplementary Figure 4. Plasma protein filtering to derive plasma signature of cognitive impairment.**

**a,** Heatmap showing results from linear models regressing each module eigenprotein with age, sex, cohort, *APOE4* dose, and cognitive impairment stage (CDR-Global). Upper heatmap shows beta effect sizes, and lower heatmap shows –log10 p-values.

**b,** Distribution of effect sizes from sex and cohort covariates from the linear models in **a**. Proteins from modules with large cohort effects (>0.25) from at least one of the cohort covariates were filtered out (left). An example protein with a large cohort effect in module 1 is shown (right).

**c,** Proteins from the module (M9) with a large *APOE* effect (as shown in **a**) were filtered out. Two example proteins from M9 with large *APOE* effects are shown.

**d,** Remaining modules were assessed for their enrichment of synapse proteins that change with cognitive impairment independent of Aβ and tau in the CSF.
